# Precision Cas9 Genome Editing *in vivo* with All-in-one, Self-targeting AAV Vectors

**DOI:** 10.1101/2020.10.09.333997

**Authors:** Raed Ibraheim, Phillip W. L. Tai, Aamir Mir, Nida Javeed, Jiaming Wang, Tomás Rodríguez, Samantha Nelson, Eraj Khokhar, Esther Mintzer, Stacy Maitland, Yueying Cao, Emmanouela Tsagkaraki, Scot A. Wolfe, Dan Wang, Athma A. Pai, Wen Xue, Guangping Gao, Erik J. Sontheimer

## Abstract

Adeno-associated virus (AAV) vectors are important delivery platforms for therapeutic genome editing but are severely constrained by cargo limits, especially for large effectors like Cas9s. Simultaneous delivery of multiple vectors can limit dose and efficacy and increase safety risks. The use of compact effectors has enabled single-AAV delivery of Cas9s with 1-3 guides for edits that use end-joining repair pathways, but many precise edits that correct disease-causing mutations *in vivo* require homology-directed repair (HDR) templates. Here, we describe single-vector, ∼4.8-kb AAV platforms that express Nme2Cas9 and either two sgRNAs to produce segmental deletions, or a single sgRNA with an HDR template. We also examine the utility of Nme2Cas9 target sites in the vector for self-inactivation. We demonstrate that these platforms can effectively treat two disease models [type I hereditary tyrosinemia (HT-I) and mucopolysaccharidosis type I (MPS-I)] in mice. These results will enable single-vector AAVs to achieve diverse therapeutic genome editing outcomes.

## Introduction

Bacterial and archaeal adaptive immune systems known as clustered, regularly interspaced, short, palindromic repeats (CRISPR), along with their CRISPR-associated (Cas) proteins, have been at the center of the RNA-guided genome engineering revolution^1,2^, including genome editing^3–11^. The most widely adopted CRISPR-Cas genome editing platforms^12,13^ are the Cas9 endonucleases that create double-strand breaks (DSBs) in target DNA sequences complementary to a designed single-guide RNA (sgRNA). In addition to the sgRNA, Cas9 requires the presence of a protospacer adjacent motif (PAM), which is a specific sequence proximal to the sgRNA-complementary target sequence^1,2,14,15^. Once cleaved, the cell relies on the DNA repair machinery to resolve these DSBs through non-homologous end joining (NHEJ), microhomology-mediated end joining (MMEJ), or homology-directed repair (HDR), the latter of which can enable a wide range of precisely determined repair outcomes^16^. More recently, base editing^17^ and prime editing^18^ have emerged as promising technologies for achieving DSB-independent precision genome editing^13^.

The most widely used Cas9 homologue has been *Streptococcus pyogenes* Cas9 [SpyCas9, 1,368 amino acids (aa)]^1,19^, which exhibits robust editing efficiencies, as well as a broad targeting range due to its short 5’-NGG-3’ PAM^1^. Additional targeting events have been enabled by SpyCas9 engineering to alter or reduce PAM requirements, and by identifying additional Cas9 homologs that are also active in eukaryotic cells but require distinct PAMs^13^. Some Cas9 homologs, including Sth1Cas9 (1,120 aa)^3^, Nme1Cas9 (1,082 aa)^20,21^, SauCas9 (1,053 aa)^22^, CjeCas9 (984 aa)^23^, Nme2Cas9 (1,082 aa)^24^, and SauriCas9 (1,061 aa)^25^ are substantially smaller. The reduced sizes are important for some modes of *in vivo* delivery, especially viral vectors such as AAV, which has great potential as a delivery modality but is severely constrained in cargo size (<5 kb in most cases)^26^. Delivery of most of these compact Cas9s within single-vector AAVs, in which both the Cas9 and its sgRNA are packaged into the same virion, has been validated *in vivo*^22–24,27,28^. AAV vectors encoding SauCas9 along with two sgRNAs^29,30^, as well as CjeCas9 with three sgRNAs^31^, have also been reported, enabling the introduction of multiplexed edits or segmental deletions. A few of the smaller Cas9s are also less prone to off-target cellular editing than wild-type SpyCas9^23,24,27,32,33^, even with guides that support efficient on-target editing, likely because of reduced enzymatic efficiencies that disproportionately dampen activity at near-cognate sites^34^.

Despite the positive attributes conferred by Sth1Cas9, Nme1Cas9, SauCas9, and CjeCas9, they suffer from a more restricted targeting range than what is imparted by SpyCas9, due to their more complex PAMs that are less abundant throughout the genome. Engineering of the PAM-interacting domain of SauCas9 reduced its wildtype PAM requirements (5’-NNGRRT-3’)^22^ to 5’-NNNRRT^35^, helping to expand its targeting flexibility. More recently, Nme2Cas9 (5’-NNNNCC-3’ PAM)^24^ and SauriCas9 (5’-NNGG-3’ PAM)^25^ were discovered as natural compact Cas9s with dinucleotide PAMs. To our knowledge, the *in vivo* utility of SauriCas9 in all-in-one AAV vectors is yet to be documented.

To date, *in vivo* genome editing applications by recombinant AAV (rAAV)-encoded Cas9 have mostly been limited to gene knockouts or insertions by cleavage events that are repaired by NHEJ pathways. Precise editing that introduces specific sequence changes that are not compatible with mismatch-mediated end joining (MMEJ) repair^36,37^ are very difficult to attain via HDR *in vivo* due in part to the requirement for co-delivery of a repair template, in addition to the low or absent activity of the HDR pathway in non-cycling cells, or during the G1 phase of the cell cycle^16^. Co-delivery of multiple AAV vectors for HDR-based precise editing *in vivo*, e.g. with the Cas9 effector on one vector, the donor on another, and the sgRNA(s) encoded by either or both vectors, has shown some promise in animal models^38–40^. Other strategies for precise editing *in vivo*, such as base editing and prime editing, involve Cas9-fused effectors that are sufficiently large as to again require two vectors^41–45^. However, the combined AAV dosages required for multi-vector strategies raise significant concerns about safety during clinical translation^46,47^, especially in light of recent events in human trials of a high-dose AAV gene therapy^48^. A recent report described a single-AAV system in which SauCas9, two sgRNAs, and an HDR donor were designed into a single >5 kb vector construct^49^. However, this study used an AAV8 capsid that lacks the VP2 subunit, yielding low vector titers with the increased cargo size. Precise genome editing strategies compatible with single-vector AAVs using well-established capsids that can be produced at pre-clinical and clinical scales are therefore still needed.

By building upon the compact size, high cellular editing accuracy, and targeting flexibility of Nme2Cas9^24,50^, we have systematically engineered novel all-in-one AAV:Nme2Cas9 vector systems with increased utility beyond that afforded by our initial version^24^, which encoded the effector and just one sgRNA. First, we showed that minimizing the Nme2Cas9 cassette, its regulatory elements, and the sgRNA provided additional coding capacity that could be used to incorporate a second sgRNA. Specific arrangements and orientations of the dual-sgRNA system were validated for their capacity to be packaged into a single AAV vector, and for their *in vivo* efficiencies in introducing two DSBs. This strategy successfully induced exon excision or inversion *in vivo* to rescue the disease phenotype of type I hereditary tyrosinemia (HT-I) mice. We then demonstrated that one of the two sgRNA cassettes could be replaced by an HDR donor, while remaining below the 5 kb cargo size required for efficient packaging into standard serotypes. We further endowed this system with Nme2Cas9 target sites flanking the HDR donor to allow self-inactivation of the vector as a function of editing effector accumulation. To enable vector packaging of a self-targeting nuclease system, we employed an anti-CRISPR (Acr) protein^51,52^ to prevent self-cleavage during vector cloning and packaging. The *in vivo* efficiency of this all-in-one HDR-based Nme2Cas9 editing system was validated in two mouse models of inherited disease [the metabolic disorder HT-I and the lysosomal storage disease mucopolysaccharidosis type I (MPS-I)]. In both cases, precise editing efficiencies reached levels sufficient for therapeutic benefit. The establishment of a self-inactivating single-rAAV system capable of precise, versatile, HDR-based genome sequence changes *in vivo*, combined with the accuracy and wide targeting range of Nme2Cas9, provide important new capabilities for the development of more potent therapeutic genome editing platforms.

## Results

### Development of a gene-editing rAAV construct with a minimized Nme2Cas9 payload

We previously engineered ∼4.7-kb rAAV gene editing vectors that deliver Nme1Cas9 or Nme2Cas9 and a 145 nucleotide (nt) sgRNA^24,27^. Both Cas9 transgenes were fused with four nuclear localization signals (NLSs) and a triple-HA epitope tag. To engineer a single Nme2Cas9 vector with additional capabilities, we created a shortened Nme2Cas9 + sgRNA backbone. Directed by Nme2Cas9 structural information^53^ as well as empirical testing of guide requirements, we truncated the repeat:anti-repeat duplex to create two shortened 121-nt and 100-nt sgRNAs (Nme.sgRNA-121 and Nme.sgRNA-100, respectively) (Supplementary Fig. 1). The truncated guides were cloned into the previously published Nme2Cas9 AAV backbone^24^. Comparison of editing efficacy at the *CYBB, LOC100505797*, and *Fah* genomic loci by plasmid transfection into HEK293T (human) and Neuro2a (mouse) cells showed that Nme.sgRNA-100 partially compromises Nme2Cas9 activity, while Nme.sgRNA-121 performance is more consistent and comparable to that of full-length Nme.sgRNA (Supplementary Fig. 1). Although Nme.sgRNA-100 performance was less consistent by plasmid transfection when driven by a U6 promoter, *ex vivo* editing by Nme2Cas9 following ribonucleoprotein (RNP) delivery using a chemically synthesized Nme.sgRNA-100 was robust. We systematically compared editing efficiencies using full-length and 100-nt sgRNAs that were made by T7 *in vitro* transcription (IVT), with and without calf intestinal alkaline phosphatase (CIP) treatment^54^, and commercial synthetic guides carrying three 2’-*O*-methyl phosphorothioate modifications at each end to protect against exonucleases^55^. Our editing data in HEK293T cells and the difficult-to-transfect T lymphoblast cell line MOLT-3 showed that the truncated Nme.sgRNA-100 conferred a higher editing efficiency compared to the other sgRNA formats when delivered as electroporated RNPs (Supplementary Fig. 1). Shortening to 100 nt places Nme2Cas9 sgRNAs within the size range needed for facile chemical synthesis for RNP delivery.

Because the 121-nt sgRNA version was more consistent as an intracellular transcript, we used it to create a shortened AAV:Nme2Cas9 vector. Using Nme.sgRNA-121 (henceforth referred to as sgRNA), a short polyadenylation signal^56^, and an Nme2Cas9 open reading frame (ORF) with two NLSs and no epitope tags, we generated a minimized AAV:Nme2Cas9:sgRNA vector with a total genome size of ∼4.4 kb (Supplementary Fig. 1; Supplementary Note). We used plasmid transfection into HEK293T and Neuro2a cells to compare the editing activity of the shortened construct to that of the previously published 4.7-kb AAV:Nme2Cas9:sgRNA backbone. Sequencing analysis indicated that the shortened AAV backbone is functional in cells with comparable editing efficiencies at three of the four sites tested (Supplementary Fig. 1). Overall, our results confirmed that the shortened AAV:Nme2Cas9 construct is efficient at genome editing, affording additional cargo capacity to expand functionality.

### Design of dual-sgRNA AAV:Nme2Cas9 vectors

Previous reports have shown that when two simultaneous DSBs occur, the NHEJ pathway can re-ligate the free DNA ends to produce a segmental deletion^30,57–60^. The delivery of two guides within the same AAV vector can support this form of editing, facilitate the simultaneous knockout of two distinct targets, or enable the use of paired nickases via single-rAAV delivery^29^. However, as with strong secondary structural elements in general^61,62^, the inclusion of two sgRNA cassettes in the same backbone can present significant obstacles to such designs by inducing the formation of truncated vector genomes^63^. To address such events, we engineered AAV:Nme2Cas9 constructs carrying two sgRNA-121 cassettes in various positions and orientations (Fig. 1a). Each of these four backbones was packaged into AAV8 capsid and the viral DNA was isolated^64^. Size comparisons using alkaline agarose gel electrophoresis analysis revealed that depending on the sgRNA-121 cassette orientation and position, the genomes of certain dual-sgRNA AAV vectors indeed produced multiple truncated forms. Two designs (henceforth referred to as Dual-sgRNA:Designs 1 and 4; Supplementary Note) appear to contain predominantly full-length, non-truncated ∼4.8-Kb genomes (Fig. 1b).

**Fig. 1.**
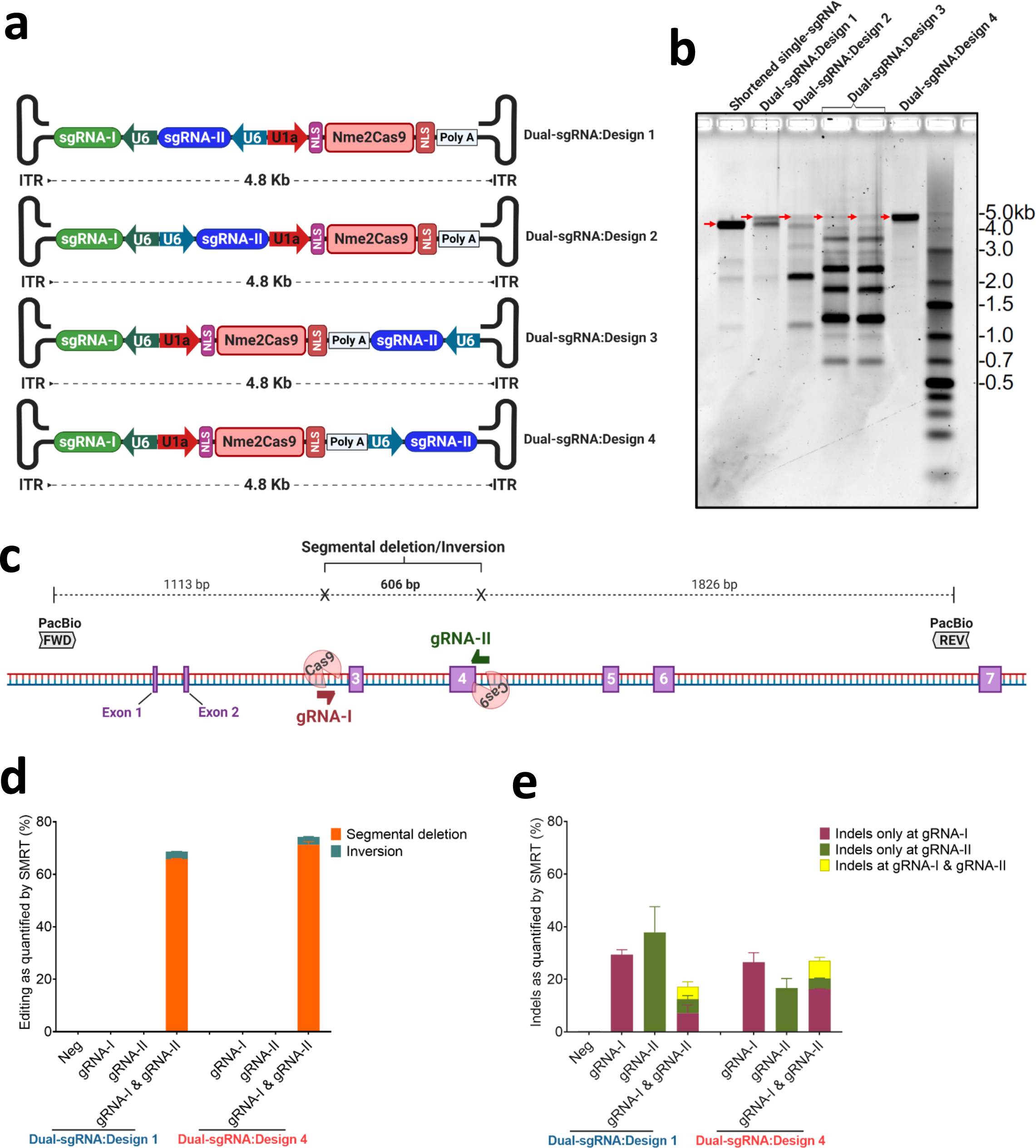
*In vitro* validation of dual-sgRNA AAV:Nme2Cas9 plasmids. **a**, Schematic of four different designs of dual-sgRNA AAV:Nme2Cas9 vector plasmids containing two sgRNA expression cassettes. **b**, Alkaline agarose gel electrophoresis of viral DNA isolated from the four dual-sgRNA AAV:Nme2Cas9 vectors presented in (**a**) after packaging in AAV8. DNA size markers are indicated. Red arrows indicate full-length non-truncated viral genomes. **c**, Schematic diagram of the mouse *Hpd* gene. The first target site is represented by half arrowhead in red (gRNA-I) and the second target site is represented in green (gRNA-II). SMRT sequencing primers are also highlighted in gray. **d**, Stacked histograms displaying the percentages of segmental deletion (orange) and inversion (blue) outcomes following gene editing after plasmid transfection of dual-sgRNA:Designs 1 and 4 plasmids in Neuro2a cells, as measured by SMRT sequencing analysis. **e**, Stacked histograms displaying the percentages of indels detected in full-length SMRT sequencing reads after *Hpd* editing by plasmid transfection in Neuro2a cells. The graph indicates genome reads with indels recorded only at gRNA-I (in red), indels only at gRNA-II in (dark green), and indels at both gRNA-I and gRNA-II (yellow) as measured by SMRT sequence analysis. Data are presented as mean values ± s.e.m. (n = 2 biological replicates).

To test the efficacy of these constructs for the induction of segmental deletion, we tested Design 1 and Design 4 for their capacity to excise a 606-bp fragment spanning exons 3 and 4 of the *Hpd* gene (Fig. 1c). We did this first in cultured cells and then in an HT-I mouse model in which the *Fah* gene, which encodes the liver enzyme fumarylacetoacetate hydrolase (FAH), is disrupted by the insertion of a neomycin cassette in exon 5 (*Fah*^*neo/neo*^)^65^. A hallmark of this disease in mice is the gradual loss of body weight when the mice are not maintained on water supplemented with the drug 2-(2-nitro-4-trifluoromethylbenzoyl)-1,3-cyclohexanedione (NTBC), which blocks the tyrosine catabolism pathway upstream of FAH, preventing the buildup of the hepatotoxic metabolites that result from loss of FAH^66^. Knocking out HPD can rescue the lethal phenotypes of HT-I mice by blocking the tyrosine catabolic pathway upstream of FAH^27,67^. In addition to the expected segmental DNA deletion, we also anticipated detecting inversions and small indels at one or both of the two target sites^30,60,68^. Initially we transfected Dual-sgRNA:Design 1 and Dual-sgRNA:Design 4 plasmids into mouse Neuro2a cells and used single molecule, real-time (SMRT) sequencing to assess editing efficiency, using primers that included unique molecular identifiers (UMIs) to enable correction for amplification bias with products of different lengths. Our analysis showed that both of these plasmids efficiently yielded the expected segmental deletion only when spacers gRNA-I and gRNA-II are co-expressed. Our data also revealed that localized, small insertions/deletions (indels) are generated at both target sites when segmental deletion does not occur (Fig. 1d,e and Supplementary Fig. 2). In addition to confirming that co-expression of Nme2Cas9 and two sgRNAs can efficiently induce segmental deletions in cultured cells^24^, these results also affirm that dual-sgRNA AAV backbones should be carefully evaluated to avoid formation of truncated genomes that may compromise vector performance *in vivo*^*63*^.

### Validation of dual-sgRNA AAV:Nme2Cas9 gene editing *in vivo*

To test the ability of dual-sgRNA AAV:Nme2Cas9 vectors to disrupt *Hpd* in HT-I mice, we used the two validated AAV8:Nme2Cas9 Dual-sgRNA:Design 1 and 4 vectors (Fig. 1b) to induce segmental deletion of exons 3 and 4 *in vivo* in *Fah*^*neo/neo*^ mice. Adults were injected with 4 x 10^11^ gc of each vector via tail vein. To assess baseline editing efficiency without selective clonal expansion of edited hepatocytes, we evaluated editing in one mouse from each group prior to NTBC withdrawal, as well as in C57BL/6 mice (Fig. 2a). After NTBC withdrawal, *Fah*^*neo/neo*^ mice were effectively rescued of the lethal phenotypes of HT-I as indicated by body weight stability for 3 weeks (Fig. 2b). In contrast, PBS-injected *Fah*^*neo/neo*^ mice lost over 10% of their body weights within 4-13 days of NTBC withdrawal and were euthanized (Fig. 2b).

**Fig. 2.**
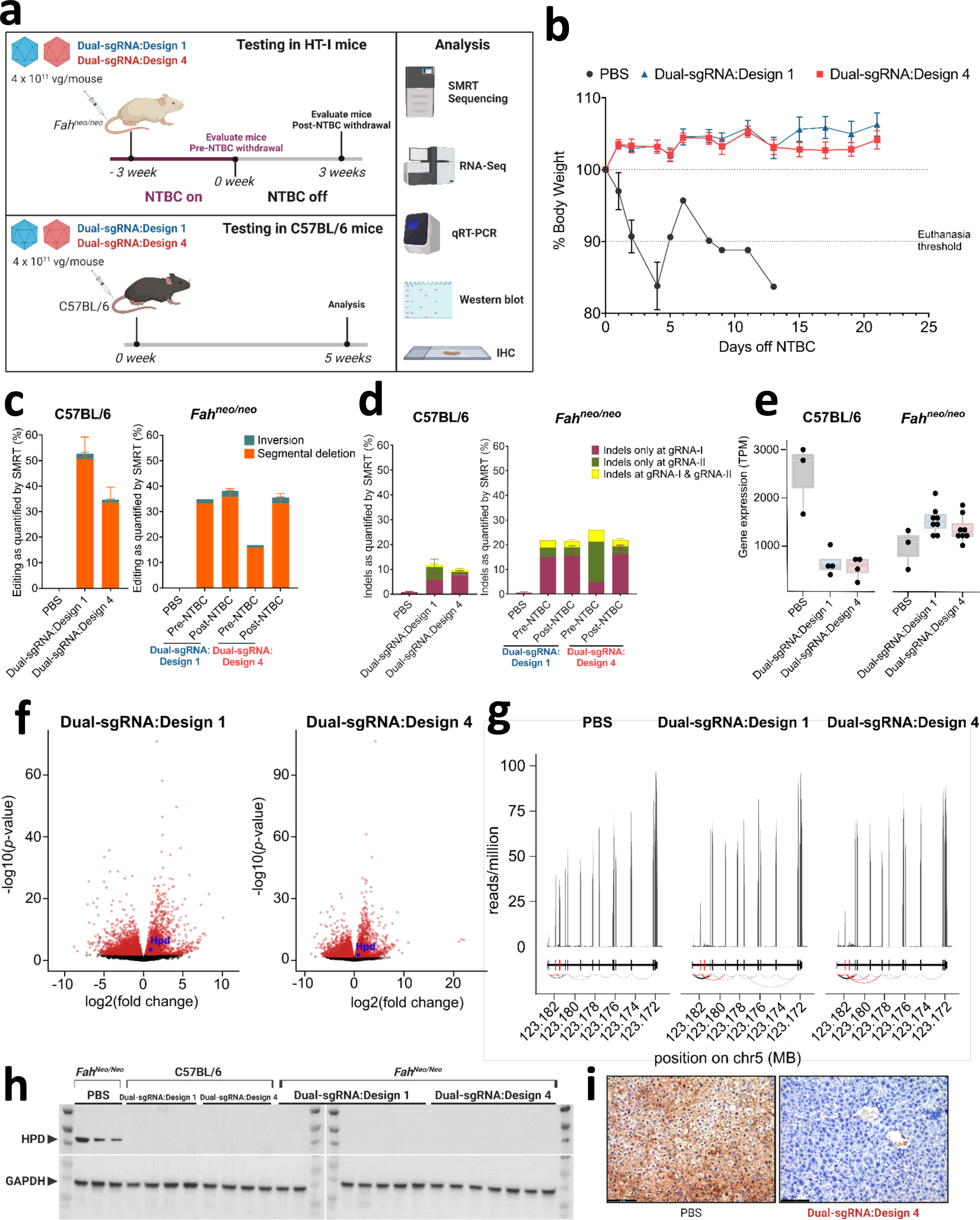
*In vivo* gene editing using dual-sgRNA rAAV:Nme2Cas9 vectors. **a**, *In vivo* experimental plan to validate rAAV:Nme2Cas9 Dual-sgRNA:Designs 1 and 4 to disrupt the *Hpd* gene by tail vein injections of AAV8 vectors in adult *Fah*^*neo/neo*^ and C57BL/6 mice. **b**, Complete rescue of body weights after Dual-sgRNA:Designs 1 and 4 in *Fah*^*neo/neo*^ mice. **c**, Quantification of editing events in C57BL/6 (left) and *Fah*^*neo/neo*^ (right) mice by SMRT sequencing analysis showing efficiencies of segmental deletion (orange) and inversion (blue) outcomes. **d**, Stacked histogram for the percentages of indels recorded in full-length (3.6 kb), UMI-corrected SMRT reads after *Hpd* editing by AAV8 delivery of Dual-sgRNA:Designs 1 and 4 in C57BL/6 (left) and *Fah*^*neo/neo*^ (right) mice. The plot indicates indels recorded only at gRNA-I (in red), indels only at gRNA-II (in dark green), and indels at both gRNA-I and gRNA-II (yellow) as measured by SMRT sequence analysis. **e**, mRNA levels from RNA-seq (transcripts per million) of *Hpd* wild-type mRNA in the livers of C57BL/6 mice (left) and *Fah*^*neo/neo*^ mice (right). **f**, Volcano plots showing differentially expressed genes (red, FDR <= 5%) between dual-sgRNA:Design 1 and PBS treated *Fah*^*neo/neo*^ mice (left) and Dual-sgRNA:Design 4 and PBS-treated *Fah*^*neo/neo*^ mice (right). **g**, RNA-seq normalized read coverage across *Hpd* exons shows a reduction of exon 4 expression in Dual-sgRNA:Design 1 and Design 4 treated *Fah*^*neo/neo*^ mice and an increase in exon-exon junction reads skipping exons 3 and/or 4. **h**, Total HPD protein knockout as shown by anti-HPD Western blot using total protein collected from mouse liver homogenates. **i**, Representative images of immunostaining for HPD in liver tissues in *Fah*^*neo/neo*^ mice injected with PBS (left) or Dual-sgRNA:Design 4 (right). Scale bar is 20 mm. Data are presented as mean values ± s.e.m. (n = 4 in C57BL/6 cohort; n = 1 in pre-NTBC *Fah*^*neo/neo*^ cohort; n = 7 in post-NTBC *Fah*^*neo/neo*^ cohort).

After completion of editing regimens, we harvested liver DNA and performed UMI-augmented SMRT sequencing of target-spanning amplicons for all mice in the study to assess the efficiencies of segmental deletion and other outcomes. These analyses revealed that segmental deletions were successfully induced by Nme2Cas9 *in viv*o with both dual-sgRNA vector designs (Fig. 2c), with the expected 3.0 kb read lengths resulting from deletion of the 606-bp fragment from the 3.6-kb amplicon (Supplementary Fig. 2). In C57BL/6 mice, which are not expected to exhibit selective proliferation of edited cells, treatment with dual-sgRNA:Designs 1 or 4 produced 50.5 ± 8.7% and 33.6 ± 6% segmental deletions between the two target sites, respectively (Fig. 2c). With the *Fah*^*neo/neo*^ cohorts treated with each vector design, the mice analyzed before NTBC withdrawal exhibited 33.3% (Design 1) and 15.8% (Design 4) segmental deletion efficiencies. Three weeks following NTBC withdrawal and selective hepatocyte expansion, vector designs 1 and 4 yielded 35.7 ± 3.3% and 33.2 ± 4% segmental deletions, respectively. Surprisingly, inversions were detected at lower frequencies than previously demonstrated (Fig. 2c)^30,60^. In addition to segmental deletions, significant frequencies of small indels were observed at the two target sites (Fig. 2d). Vector fragment integration at the DSB sites were detected at a frequency of 2.8 ± 0.7% (Design 1) and 2.4 ± 0.8% (Design 4) in C57BL/6 mice, at 2.3% (Design 1) and 1.5% (Design 4) before NTBC withdrawal, and 3.5 ± 0.5% (Design 1) and 2.7 ± 0.5% (Design 4) post-NTBC withdrawal (Supplementary Fig. 2).

We extensively characterized genome-wide mRNA levels using mRNA extracted from mouse livers by mRNA sequencing. Our data revealed that efficient segmental deletion in recovered *Fah*^*neo/neo*^ cohorts treated with each dual-sgRNA vector design yielded *Hpd* mRNA levels that were lower than those observed in healthy, PBS-treated wild-type mice (Figs. 2e and 2f). *Hpd* mRNA levels were also strongly reduced (relative to PBS-treated C57BL/6 mice) in PBS-injected *Fah*^*neo/neo*^ animals, presumably as a secondary consequence of extensive liver damage. *Hpd* mRNA levels were significantly decreased in C57BL/6 mice treated with either of the dual-sgRNA vectors (Fig. 2e), reflecting the expected results of Nme2Cas9 editing. Globally, 4,792 and 5,027 genes were significantly differentially expressed in *Fah*^*neo/neo*^ mice after treatment with Dual-sgRNA:Design 1 and 4 vectors, respectively (FDR <= 5%, relative to PBS treatment). Differentially expressed genes were enriched within similar gene ontology categories across the two dual-sgRNA treatments, with an enrichment of genes in metabolic processes, regulation of endopeptidase activity, and response to external stimulus (Supplementary Table). In both cohorts of mice, there was a decrease in the inclusion of exons 3 and 4 in mature *Hpd* mRNA transcripts (Fig. 2g), with both exons 3 and 4 being skipped in 4.2-5.5% of transcripts (compared to 0.08% of transcripts in PBS treated mice). More strikingly, due to the proximity of gRNA-II to the splice donor site (Fig. 1c), exon 4 is excluded in 86.7-87.5% of *Hpd* mRNAs in dual-sgRNA treated mice (compared to 0.33% of transcripts in PBS treated mice) (Supplementary Fig. 2). Consistent with the observed prevalence of segmental deletions, inversions, skipped exons and other perturbations (Fig. 2c,d), our protein analyses showed efficient knockdown of HPD in the livers of the edited *Fah*^*neo/neo*^ and C57BL/6 mice by Western blot (Fig. 2f) and immunohistochemistry (Fig. 2g; Supplementary Fig. 2). These data demonstrate that Dual-sgRNA:Designs 1 and 4 are effective vectors for inducing segmental DNA deletions *in vivo*.

### Design of self-eliminating rAAV:HDR vectors

The majority of rAAV-based applications to repair pathogenic mutations *in vivo* rely on a two-vector system for the delivery of Cas9 in one rAAV and the HDR donor (along with the guide) in another^38,69,70^. Although this strategy allows control over dosing at different donor:effector ratios, it also requires higher overall vector doses, which might be toxic to the patient. Furthermore, the lack of simultaneous transduction with both vectors in a percentage of cells will limit editing efficiencies. Previously, HDR-based precision editing platforms involving simultaneous delivery of Cas9, sgRNA, and donor DNA in a single vehicle generally used adenoviral vector^71^ or RNP^72,73^ delivery, neither of which can reach all cell types of clinical interest. To extend single-vehicle HDR-based gene editing to AAV vectors that are within the standard limits of efficient packaging (∼4.9 kb) with well-characterized capsids, we engineered rAAV backbones containing Nme2Cas9, sgRNA and a <500-nt donor DNA for *in vivo* testing of HDR editing.

To examine the best placement of the donor DNA within the rAAV backbone, we first created two designs in which the donor was placed near the 3’ ITR (design A) or in the middle of the backbone (design B) (Fig. 3a). We also created designs C and D, which respectively differ from A and B only in the presence of Nme2Cas9 sgRNA target sites flanking the donor (Fig. 3a). The presence of Cas9 target sites that flank an HDR donor has previously been shown to increase HDR efficiencies in cell culture after plasmid transfection^74^. These rAAV:HDR:cleaved designs also have the potential to enable self-elimination of the AAV:Nme2Cas9 vector, limiting or preventing long-term Cas9 expression *in vivo*^75–77^ that could exacerbate off-target mutagenesis, genotoxicity, and immunogenicity.

**Fig. 3.**
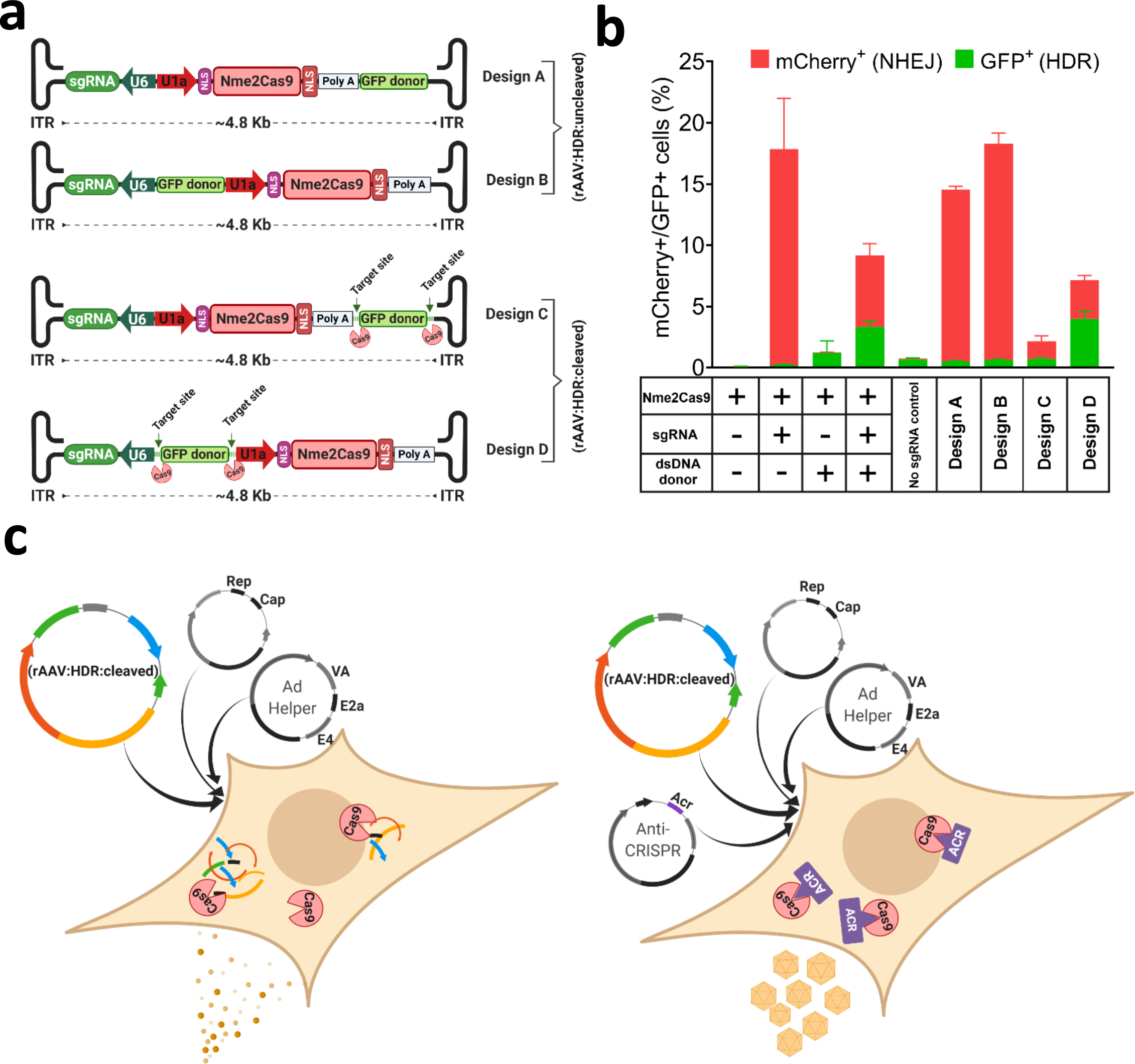
*In vitro* design and validation of all-in-one rAAV:HDR vectors. **a**, Schematic of four different vector designs of rAAV:HDR constructs containing an sgRNA expression cassette, Nme2Cas9, and a donor DNA (<500 bp) with and without flanking sgRNA target sites. **b**, Efficiencies of NHEJ and HDR events depicted as percentages of mCherry- and GFP-positive cells, respectively, obtained after transfection of AAV:Nme2Cas9:sgRNA and dsDNA GFP donor *in trans*), or of rAAV:HDR constructs in TLR-MCV1 HEK293T cells. Data are presented as mean values ± s.e.m. (n = 3 biological replicates). **c**, Schematic of modified rAAV packaging transfection system using anti-CRISPR protein plasmid to block self-targeting by Nme2Cas9 expression in packaging cells during production.

The major hurdle in creating the rAAV:HDR:cleaved design C and D backbones was the degradation of the plasmids after transformation in *E. coli* competent cells due to leaky expression of Nme2Cas9 and sgRNA in bacteria. To resolve this issue, we used λred phage recombineering technology^78^ to integrate the *H. parainfluenzae* anti-CRISPR *acrIIc4 Hpa* gene^79^, under the control of its native promoter, into DH5α competent *E. coli* cells to inhibit Nme2Cas9 and prevent plasmid degradation. These plasmids were tested in a Traffic Light Reporter (TLR) system^80^, TLR-Multi-Cas-Variant 1 (MCV1), that has a disrupted GFP coding sequence followed by an out-of-frame mCherry cassette^81^. The reporter cassette was inserted as a single copy via lentiviral transduction in HEK293T cells. When a nuclease cleaves the insert that disrupts *eGFP*, some NHEJ-induced indels place the mCherry in frame, inducing red fluorescence. By contrast, if an *eGFP* HDR donor is supplied and used by the cell, the disrupted *eGFP* is repaired, inducing green fluorescence. We cloned an *eGFP* donor and an *eGFP*-insert-specific Nme2Cas9 sgRNA into the rAAV:HDR constructs to enable HDR to restore eGFP expression; in designs C and D, we also included sgRNA target sites flanking the donor. Our flow cytometry analyses show that design D resulted in eGFP signals that were comparable to those induced by the use of a separate linear donor (generated by PCR) (Fig. 3b). We chose design D and its non-self-inactivating analog (design B) (Supplementary Note) for *in vivo* testing and validation.

### *In vivo* validation of rAAV:HDR vectors in HT-I mice

To examine the therapeutic potential of our all-in-one rAAV:HDR systems, we designed two vectors: rAAV:HDR:uncleaved (from design B) and the self-eliminating rAAV:HDR:cleaved (from design D) (Fig. 3a). For these studies we used HT-I mice that contain a point mutation in the last nt of exon 8 of the *Fah* gene^66^ (*Fah*^*PM/PM*^), leading to exon skipping that disables expression of the FAH protein^82^. Both vector designs contain Nme2Cas9, sgRNA targeting *Fah*, and a 358bp HDR donor (including homology arms) to correct the G-to-A point mutation in *Fah* exon 8 in HT-I *Fah*^*PM/PM*^ mice. The HDR donor sequence also included PAM mutations (CC > TT, within intron 8) at the target site to prevent cleavage of the donor fragment or of the HDR-repaired *Fah* locus. In addition, we created a negative control vector expressing an sgRNA with a non-cognate spacer sequence (rAAV:FahDonor:ncSpacer) and another with a non-cognate donor (rAAV:ncDonor:FahSpacer) (Fig. 4a). Neither of these negative control vectors is capable of self-targeting. These four ∼4.8-kb vectors were packaged by the triple transfection method for rAAV8 packaging^64^. Packaging of the rAAV:HDR:cleaved vector was modified to include a fourth plasmid expressing AcrIIC4*Hpa* to prevent the self-elimination of the rAAV:HDR:cleaved plasmid during production (Fig. 3c).

**Fig. 4.**
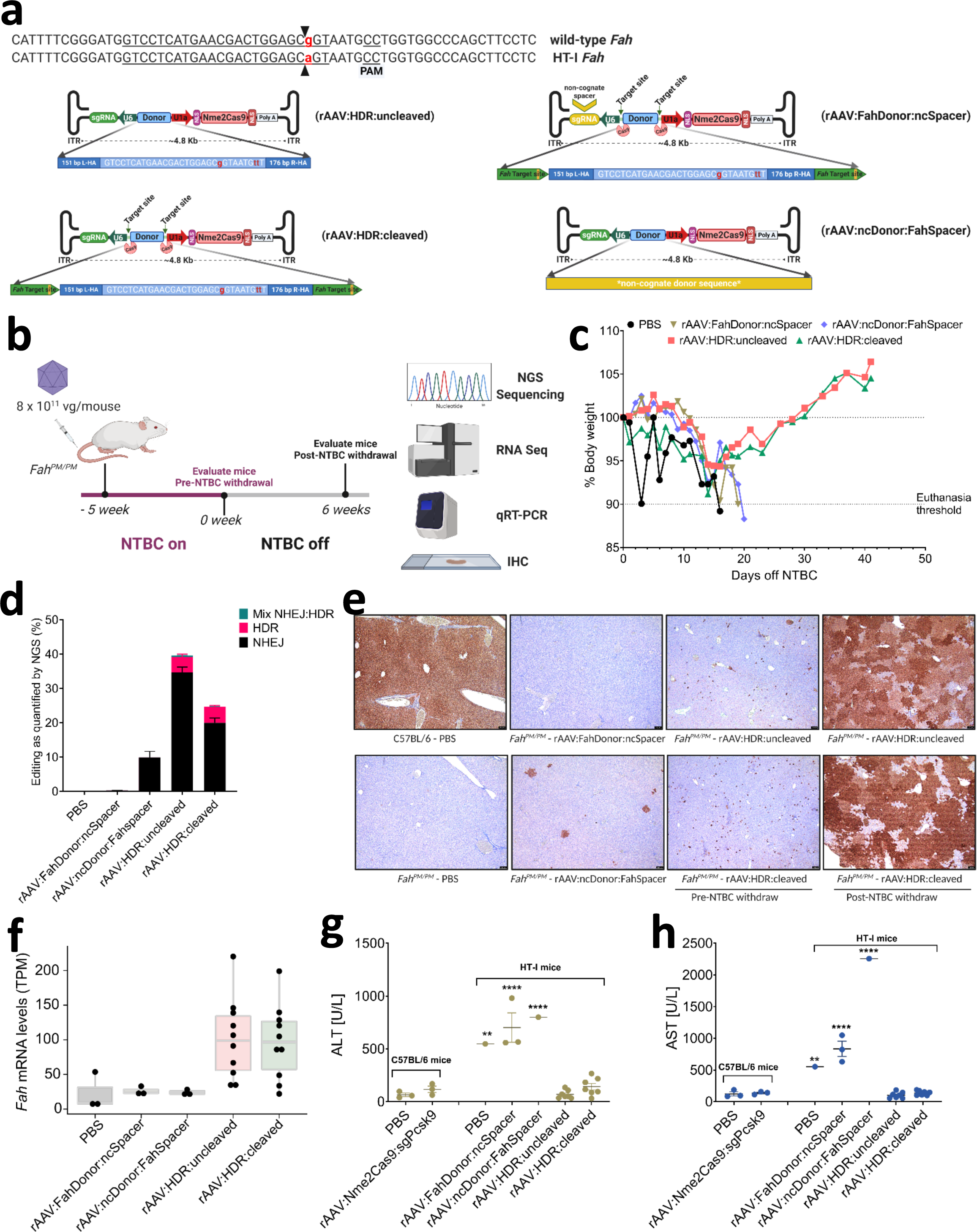
rAAV:HDR vectors rescue liver disease phenotypes in a mouse model of HT-I. **a**, Schematic of rAAV:HDR:uncleaved and -cleaved vectors (left) and control non-cognate donor or spacer vectors (right) to correct the *Fah* point mutation in *Fah*^*PM/PM*^ mice (top). **b**, *In vivo* experimental regimen to correct the *Fah* mutation by injection of AAV8 though the tail vein in adult *Fah*^*PM/PM*^ mice. **c**, Complete rescue of average body weight loss in *Fah*^*PM/PM*^ mice in treated cohorts injected with rAAV:HDR:uncleaved (red) and -cleaved (green), as compared to control cohorts injected with PBS (black), rAAV:FahDonor:ncSpacer (brown) or rAAV:ncDonor:FahSpacer (blue). Data shown represent the average body weight of each cohort. **d**, Stacked histograms showing the percentage distribution of NHEJ, HDR, and imprecise NHEJ:HDR mix at *Fah* in livers of mice 6 weeks after NTBC withdrawal, as measured by NGS sequencing of PCR amplicons from genomic DNA. **e**, Representative images of immunostaining for FAH in liver tissues of negative control and treated cohorts, as indicated below each panel. Scale bar, 100 mm. **f**, Levels of *Fah* mRNA from RNA-seq (transcripts per million) in the livers of *Fah*^*PM/PM*^ mice. **g**, Serum ALT activity in WT C57BL/6 mice injected with PBS and rAAV:Nme2Cas9 vector targeting *Rosa26* from a previous study, in comparison to untreated negative control and treated *Fah*^*PM/PM*^ mice. Statistical analysis used one-way ANOVA (*p* < 0.0001) with Dunnett’s test. Data are presented as mean values ± s.e.m. \*\**p* = 0.0025, \*\*\*\**p* < 0.0001 vs. PBS-injected C57BL/6 group. **h**, Serum AST activity in WT C57BL/6 mice injected with PBS and rAAV:Nme2Cas9 vector targeting *Rosa26* from a previous study, in comparison to untreated negative control and treated *Fah*^*PM/PM*^ mice. Statistical analysis used one-way ANOVA (*p*<0.0001) with Dunnett’s test. Data are presented as mean values ± s.e.m. \*\**p* = 0.0011, \*\*\*\**p* < 0.0001 vs PBS-injected C57BL/6 group. Data are presented as mean values ± s.e.m. (n = 3 in PBS, rAAV:FahDonor:ncSpacer, rAAV:ncDonor:FahSpacer and pre-NTBC withdrawal AAV.HDR:cleaved and -uncleaved cohorts; n = 7 in post-NTBC withdrawal rAAV:HDR:cleaved and -uncleaved cohorts).

For *in vivo* evaluation, 8 x 10^11^ gc of each vector was injected into adult HT-I *Fah*^*PM/PM*^ mice via tail vein. Mice were maintained on NTBC for five weeks and then monitored for body weight loss after NTBC withdrawal (Fig. 4b). Mice injected with PBS, rAAV:FahDonor:ncSpacer, or rAAV:ncDonor:FahSpacer showed the expected gradual body weight loss and were euthanized. Strikingly, cohorts injected with rAAV:HDR:uncleaved and rAAV:HDR:cleaved initially lost weight, but recovered and stayed healthy for 41 days (Fig. 4c). Some mice in the rAAV:HDR:cleaved and -uncleaved cohorts were put on NTBC-containing water temporarily to promote initial recovery.

To determine the efficiency of genome editing, we performed targeted deep sequencing analysis to measure the level of genome modification at the *Fah* locus. HDR efficiency, calculated by co-conversion of all three edited positions [A..CC to G..TT], in pre-NTBC cohorts was very low (<0.1%); accurate quantifications were difficult to attain because of the inherent error rate of NGS, as well as the polyploid nature of hepatocytes and the presence of DNA from other nonparenchymal cells that are not efficiently transduced by AAV8^82^. HDR-edited reads were significantly increased to 4.4 ± 0.9% and 4.7 ± 0.4% in the rAAV:HDR:uncleaved and rAAV:HDR:cleaved cohorts, respectively 6 weeks after NTBC withdrawal. The relative increase over apparent editing efficiency in the wildtype cohort was likely due to the selective growth advantage of the successfully edited *Fah*^*+*^ cells after NTBC withdrawal (Fig. 4d; Supplementary Fig. 3).

We also performed immunohistochemistry analyses of liver tissues to examine the extent of liver recovery by FAH protein expression. Before drug withdrawal, we detected a marked increase in the number of FAH^+^ cells in both rAAV:HDR cohorts (Fig. 4e; Supplementary 4). Interestingly, we observed through amplicon sequencing analysis that there were sporadic FAH^+^ cells in mice injected with rAAV:ncDonor:FahSpacer as a result of mutations that likely result from NHEJ repair events at the adjacent Nme2Cas9-catalyzed DSB (Supplementary Fig. 4). Additionally, those cells were likely expanded after NTBC withdrawal, as the starting population was apparently insufficient to allow the mice to recover from the disease phenotype in the absence of NTBC augmentation (Fig. 4e; Supplementary Fig. 4). A greater number of corrected FAH^+^ cells were observed in the rAAV:HDR cohorts before drug withdrawal, supporting eventual recovery (Fig. 4e; Supplementary Fig. 4). mRNA sequencing analyses showed that effective gene repair post-NTBC withdrawal was associated with significant increases in the level of *Fah* mRNA in both rAAV:HDR cohorts (relative to PBS, ncDonor, and ncSpacer controls, Fig. 4f and Supplementary Fig. 3). RT-PCR analysis with primers spanning exons 5 to 9 of *Fah* showed that the increased levels of *Fah* mRNA is also associated with exon 8 inclusion (Supplementary Fig. 3).

As expected by body weights, editing efficiencies, and FAH expression, HT-I mice treated with rAAV:HDR vectors exhibited striking reductions in liver damage as indicated by the significant reduction in serum aspartate aminotransferase (AST) and alanine aminotransferase (ALT) levels (Fig. 4g,h), as well as normalization of hepatocyte histology (Supplementary Fig. 3). Globally, 8,827 and 8,170 genes were differentially expressed in HT-I mice treated with rAAV:HDR:uncleaved and rAAV:HDR:cleaved vectors, respectively (FDR <= 5%, relative to treatment with rAAV:HDR:ncDonor:FahSpacer vector; Supplementary Fig. 3). Differentially expressed genes were enriched among gene ontology categories involved in small molecule metabolic processes (Supplementary Table). Only four genes were significantly differentially expressed between the rAAV:HDR:uncleaved and rAAV:HDR:cleaved vector treatments (FDR <= 5%, Supplementary Fig. 3), indicating similar targeting ability between the two vectors.

Previous efforts to create self-targeting rAAV:Cas9 vectors involved dual-vector systems^77,83^. Our all-in-one system necessarily involves co-expression of Nme2Cas9 and sgRNA in the same cell, enabling the targeting of engineered sites flanking the *Fah* HDR donor (Fig. 4a). Quantitative PCR was used to examine the number of rAAV genome copies in mice and suggested that liver samples from animals treated with the rAAV:HDR:cleaved vector contained significantly lower rAAV copies than the rAAV:HDR:uncleaved mouse cohorts after NTBC withdrawal (Supplementary Fig. 5). This reduction in rAAV episomal numbers was not associated with a significant decrease in steady-state levels of Nme2Cas9 mRNA as measured by mRNA sequencing (Supplementary Fig. 5). However, total Nme2Cas9 mRNA and sgRNA transcripts decreased over time, as measured by quantitative RT-PCR with random hexamer primers capturing both nascent and mature mRNAs, suggesting that the AAV:HDR:cleaved platform can reduce the unwanted long-term expression of Nme2Cas9 (Supplementary Fig. 5). Additionally, we observed that the detected level of rAAV fragments integrated at the Cas9 DSB *Fah* site is significantly lower in the rAAV:HDR:cleaved cohort compared to rAAV:HDR:uncleaved (Supplementary Fig. 5). All of these data suggest that the self-eliminating rAAV:HDR:cleaved vector is a more therapeutically viable option, as it can function *in vivo* to repair DNA mutations, while eliminating rAAV genomes and Nme2Cas9 expression over time.

### *In vivo* validation of rAAV:HDR vectors in MPS-I mice

The selective expansion of correctly edited cells in HT-I mice enabled therapeutic benefit even when editing efficiencies were very low. We sought a second disease model in which editing is not known to induce selective clonal expansion, to determine whether HDR efficiencies with our all-in-one rAAV system could still reach therapeutic thresholds. We chose MPS-I mice, which suffer from an accumulation of glycosaminoglycans (GAGs) in lysosomes. These mice carry a G-to-A mutation in the gene encoding the α-L-iduronidase (IDUA) enzyme, converting the Trp392 codon into a stop codon. Previous reports showed that IDUA enzyme activity as low as 0.2% of normal levels can provide therapeutic benefit in MPS-I patients^84^, largely due to uptake of secreted IDUA by uncorrected cells. We engineered two *Idua*-targeting vectors analogous to those used above (rAAV:HDR:uncleaved and rAAV:HDR:cleaved) containing Nme2Cas9, an sgRNA targeting *Idua*, and a 272-bp donor (including homology arms) to correct the disease-causing G-to-A point mutation. Because the pathogenic mutation resides within an exonic protein-coding sequence, a PAM mutation is not feasible without altering the amino acid sequence. As an alternative approach, we incorporated two silent mutations (A..C > G..T) in the seed sequence of the spacer to prevent cleavage of the donor (Fig. 5a). Re-targeting of the edited allele by Nme2Cas9 would be further inhibited by the third non-complementary seed residue at the HDR-corrected position. For the negative control vector, we generated an rAAV backbone analogous to rAAV:HDR:uncleaved, but containing a non-cognate sgRNA spacer (rAAV:IduaDonor:ncSpacer) (Fig. 5a). All three vectors were packaged into AAV9 capsids, and cohorts of neonate MPS-I mice were injected with 4 x 10^11^ gc of each vector via facial vein (Fig. 5b). Liver tissue lysates were collected and IDUA enzymatic activities were measured. We observed significant increases in IDUA activity to 3.0 ± 0.03% and 2.3 ± 0.2% of wild-type levels in rAAV:HDR:uncleaved and rAAV:HDR:cleaved cohorts, respectively (Fig. 5c). This increase was above the therapeutic threshold required for effective reduction of GAG accumulation in the liver (Fig. 5d). Targeted amplicon sequencing analysis of total liver DNA confirmed the corrected “G--T-G” pattern at the *Idua* locus in rAAV:HDR:uncleaved (4.5 ± 0.49%) and rAAV:HDR:cleaved (3.3 ± 0.43%) cohorts (Fig. 5e). Via qRT-PCR assays we also observed significant increases in *Idua* mRNA in the liver (Fig. 5f), likely due to reductions in nonsense-mediated mRNA decay of the edited transcripts. Other organs exhibited lower but notable levels of HDR (Fig. 5e).

**Fig. 5.**
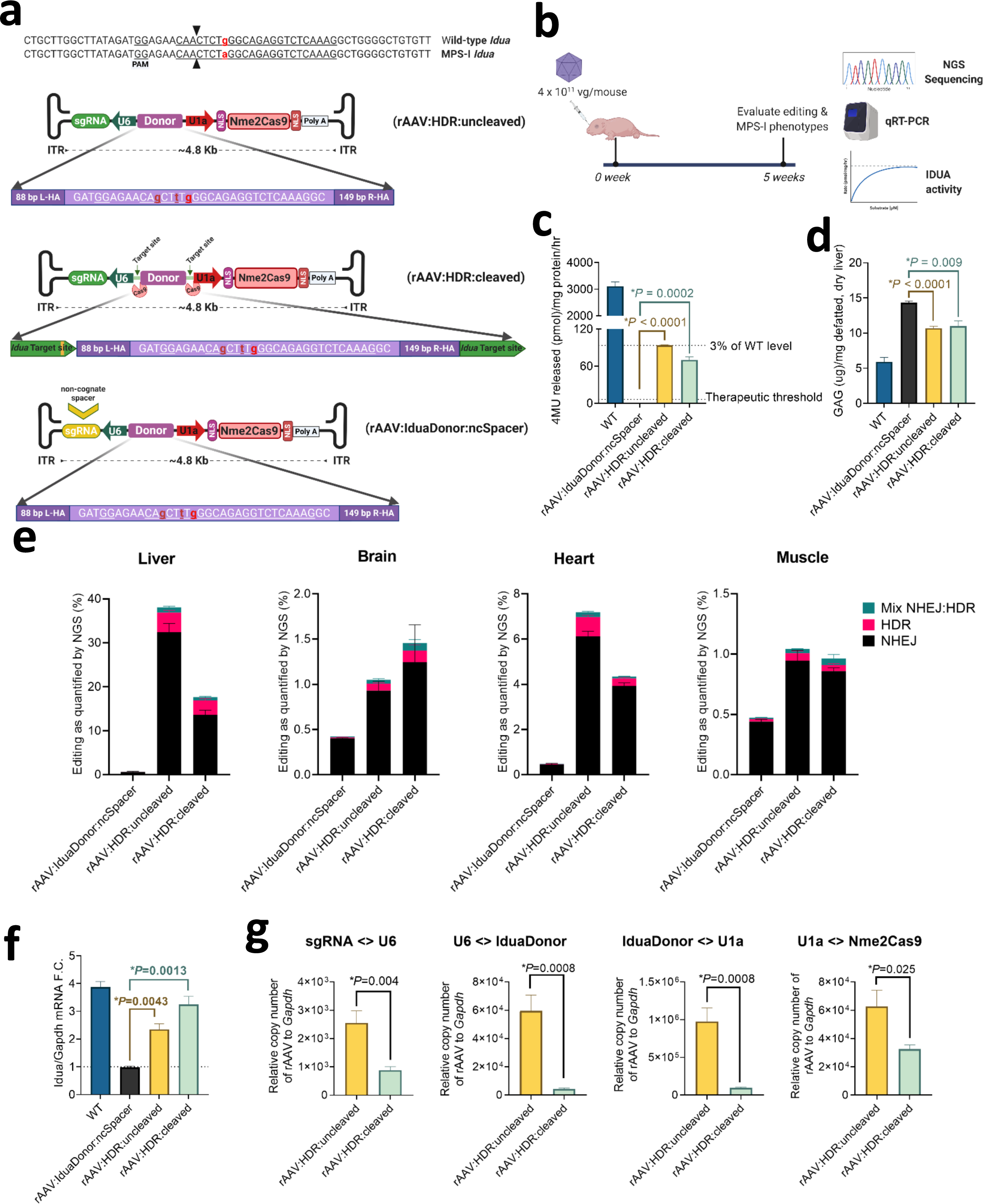
Reduced clinical disease manifestation in a mouse model of mucopolysaccharidosis type I treated with rAAV:HDR vectors. **a**, Schematic of rAAV:HDR:uncleaved, -cleaved, and control non-cognate spacer vectors to correct the *Idua* point mutation in MPS-I *Idua*^*W392X*^ mice. **b**, *In vivo* experimental regimen to correct the *Idua* mutation by injection of rAAV9 though the facial vein in neonate MPS-I mice. **c**, IDUA specific activity in liver lysates of healthy mice compared to rAAV:IduaDonor:ncSpacer mice (negative control) and treated rAAV:HDR uncleaved/cleaved cohorts. The dashed lines indicate 0.2% (therapeutic threshold) and 3% of the IDUA activity detected in lysates from WT mice. **d**, Level of glycosaminoglycan (GAG) accumulation in the livers of the same mice shown in (**c**). **e**, Stacked histogram showing the percentage distribution of NHEJ, HDR and imprecise NHEJ:HDR mix at *Idua* in liver, brain, heart and muscle tissues, as measured by NGS sequencing of PCR amplicons from genomic DNA. **f**, qRT-PCR data showing increase in *Idua* mRNA in the liver normalized to *Gapdh* mRNA. **g**, AAV copy numbers in MPS-I mouse liver tissues using the [sgRNA<>U6], [U6<>IduaDonor], [IduaDonor<>U1a] and [U1a<>Nme2Cas9] primer pairs. Data are presented as mean values ± s.e.m. (n= 4-5 biological replicates). *p* values are calculated using Student’s t-test (2-sided).

To examine the self-cleaving ability of rAAV:HDR:cleaved vectors, quantitative PCR assays were designed using four primer pairs, including two that span the ends of the donor (where the cleavage sites reside in the self-targeting version) (Supplementary Fig. 6). Unlike rAAV:HDR:uncleaved cohorts, mice injected with the rAAV:HDR:cleaved vector had significantly lower qPCR signals in the liver (Fig. 5g). The frequency of rAAV fragment integration at the *Idua* target site (as detected by NGS sequencing) was minimal, with significantly fewer insertion events detected with the rAAV:HDR:cleaved vector (Supplementary Fig. 6). Nme2Cas9 mRNA and sgRNA transcripts were modestly reduced, as measured by qRT-PCR (Supplementary Fig. 6). Since these mice were injected as neonates, Nme2Cas9-specific humoral immune responses did not develop in MPS-I mice, while adult-treated HT-I and healthy C57BL/6 mice treated with rAAV:Nme2Cas9 targeting the *Rosa26* locus^24^ generated readily detectable α-Nme2Cas9 IgG1 antibodies with no apparent difference between rAAV:HDR:cleaved and rAAV:HDR:uncleaved cohorts (Supplementary Fig. 6). In conclusion, rAAV:HDR vectors have the potential to correct inherited disease mutations *in vivo*, while enabling the therapeutic benefits of longer-term self-elimination to minimize genotoxic and off-target effects, as well as possible cellular immune responses against transduced tissues.

## Discussion

The tremendous potential of precision genome editing *in vivo* to repair pathogenic mutations still faces significant translational hurdles. AAV vectors for Cas9 genome editing exhibit great promise for some tissues and are already in clinical trials for NHEJ-based edits (ClinicalTrials.gov; identifier: NCT03872479). However, rAAV delivery systems for precise genomic re-writing via HDR or base editing have required multiple vectors^38–45^. The extreme rAAV dosages needed for multi-vector strategies impose safety difficulties in clinical applications^46–48^. Here, we have developed single-vector systems with standard rAAV serotypes and cargo limits for precise Nme2Cas9 editing via HDR. Nme2Cas9 is well-suited for this strategy because it is compact, active in adult mammals *in vivo*, highly accurate within eukaryotic cells, and has a broad targeting range due to its modest (dinucleotide) PAM requirement^24,50^. We further employed an Acr protein during cloning and vector production to enable the inclusion of self-targeting sites that can facilitate vector self-elimination over time, once the Nme2Cas9/sgRNA complex has accumulated to functional levels. The use of Acr proteins to protect otherwise self-inactivating genome editing vectors was reported previously for adenovirus vectors^85^ but to our knowledge has not been described for the more commonly employed rAAV. The ever-expanding roster of anti-CRISPRs against different genome editing effectors^51,52^ suggests that this strategy will be useful far beyond Nme2Cas9. We validated our all-in-one rAAV HDR platform *in vivo* in two disease models in mice (HT-I and MPS-I), and in both cases editing efficiencies reached therapeutic thresholds. Our ability to fit an HDR donor into a single-vector system along with Nme2Cas9-NLS and sgRNA transgenes was enabled by minimizing the latter components and their regulatory elements. We also generated all-in-one vectors encoding two guides, and demonstrated that this dual-sgRNA all-in-one system was effective *in vivo* for inducing segmental deletions in HT-I mice. In the latter case, sgRNA cassette placement and orientation was a critical factor in vector packaging efficiency. The two successful designs involved precise targeting of sgRNAs to exon boundaries, resulting in an unexpected diversification of *Hpd* mRNA splicing events through unknown mechanisms that merit further study.

Although therapeutic thresholds were reached by the all-in-one HDR system *in vivo* for the two preclinical disease models that we tested, HDR efficiencies were nonetheless low in absolute terms. This is not particularly surprising because of several factors: (*i*) the lengths of the homology arms needed for HDR are constrained by rAAV vector cargo limits; (*ii*) donor DNA cannot be delivered in excess of guide- and effector-expressing vector; (*iii*) many mammalian cells and tissues *in vivo* are quiescent or terminally differentiated, leading to a general loss of HDR capacity^16^. In its current form, our single-AAV-vector HDR approach would therefore be limited to applications where successfully edited cells have a proliferation advantage (as with HT-I), or where low levels of correction suffice for clinical benefit (as with MPS-I). Nonetheless, in such cases, the reduction in vector dose permitted by the use of a single-vector system could be crucial for the safety profile of precision therapeutic editing. A previous report also described an all-in-one rAAV system for HDR-based editing and applied it to HT-I mice^49^. However, this method used a VP2-deleted capsid system that enabled larger cargoes (>5 kb) but appeared to package inefficiently^49^. We report here that standard rAAV vector platforms that accommodate <4.9 kb genomes can be used for single-vector HDR *in vivo*.

In addition to reductions in vector dose, safety prospects are also potentially improved by our system’s implementation of self-cleavage ability, enabled by anti-CRISPR inhibition of Cas9 activity during cloning and packaging. Our qPCR assays suggest modest degrees of rAAV genome depletion over a 5-to 11-week editing regimens. Whether self-elimination increases over longer time courses will require more extended follow-up studies. We did not observe detectable increases in vector fragment insertion into the chromosomal target site with the self-inactivating vector. We designed our self-targeting system to have the added benefit of generating a linear HDR donor, in part to test the hypothesis that such a configuration would improve HDR efficiency over that of the uncleaved HDR vector. We obtained no clear evidence that donor excision capacity improves HDR frequencies *in vivo*, though the potential long-term benefits of vector elimination still apply, even if HDR enhancement via donor cleavage cannot be detected in other physiological contexts in the future.

In conclusion, we have developed and validated new viral delivery platforms for *in vivo* genome editing that enable precision editing via HDR, using single-vector systems that are compatible with well-characterized rAAV backbones and serotypes. We have further developed the means to enable cloning, packaging, and use of rAAV vectors endowed with self-targeting capabilities. These systems improve the balance between the competing imperatives of improving vector capabilities (e.g. enabling precise editing via HDR) and limiting vector dose to maximize safety, reduce cost, and mitigate production bottlenecks. As additional routes toward increased HDR emerge in the future, this could enable single-vector HDR-based rAAV delivery systems to address clinical indications that demand higher levels of precise gene correction, perhaps including those that are difficult or impossible to achieve via base editing or prime editing.

## Methods

### Hereditary Tyrosinemia type I mice

All animal study procedures were approved by the Institutional Animal Care and Use Committee (IACUC) at the University of Massachusetts Medical School. HT-I mice were kindly provided by Dr. M. Grompe and maintained on C57 background for the *Fah*^*PM/PM*^ strain, and on a 129 background for *Fah*^*neo/neo*^. Mice were kept on 10 mg/L 2-(2-nitro-4-trifluoromethylbenzoyl)-1,3-cyclohexanedione (NTBC; Sigma-Aldrich, Cat. No. PHR1731-1G) in drinking water when indicated. Male and female mice (8 to 15 weeks old) were used, and their body weights were recorded every 1-3 days. Moribund mice with more than 10% body weight loss were humanely euthanized, and sera and liver tissues were harvested for genome editing analyses. Tissues were pulverized in liquid nitrogen and aliquots of tissue powder (∼30 mg) were used to extract total DNA [using the DNeasy Blood and Tissue Kit (Qiagen), according to the manufacturer’s protocol], as well as total RNA and total protein.

### Mucopolysaccharidosis type I mice

Homozygous MPS-I *Idua*^*W392X*^ mice^86^ were purchased from the Jackson Laboratory (Stock No. 017681) and used to breed neonatal *Idua*^*W392X*^ pups for rAAV injections. Postnatal day 1 mice were injected with 4 x 10^11^ gc of rAAV:HDR:uncleaved, rAAV:HDR:cleaved, or control vectors via facial vein. Five weeks after injection, blood was collected via cardiac punch, and mice were transcardially perfused with ice-cold PBS. Tissues were immediately dissected, snap-frozen in liquid nitrogen, and stored at −80°C. Both female and male animals were used. All animal procedures were reviewed and approved by the IACUC at University of Massachusetts Medical School and performed in compliance with all relevant ethical regulations.

### Cloning of rAAV plasmids

All rAAV vectors were based on our previously published rAAV plasmid [Nme2Cas9_AAV; Addgene plasmid #119924] design^24^, with modifications to increase the cargo capacity (Supplementary Fig. 1). Human-codon-optimized *Neisseria meningitidis* DE10444 Cas9 (Nme2Cas9) is under the expression of a *U1a* promoter, with one SV40 nuclear localization signal (NLS) on the N terminus and one nucleoplasmin NLS on the C terminus, followed by a short poly(A) signal^56^. Shortened single-guide RNA Nme.sgRNA-121 was synthesized as a gblock for cloning. Target spacer sequences were inserted by digesting the plasmid with BspQI restriction enzyme, and the annealed spacer sequence carrying appropriate BspQI overhang sequences were ligated. This plasmid is available on Addgene (pEJS1089: mini-AAV:sgRNA.Nme2Cas9; Addgene # 159536).

### Cloning of Dual-guide rAAV:Nme2Cas9 plasmids

A second human *U6* promoter with an Nme.sgRNA-121 expression cassette was generated as a gblock (IDT) and cloned into the minimized rAAV:Nme2Cas9 plasmid to create dual-sgRNA designs 1-4 (Fig. 1a). The two configurations that enabled efficient packaging of full-length vector genomes (designs 1 and 4) are available on Addgene (pEJS1099: Dual-sgRNA:Design 4, Addgene #159537; pEJS1096: Dual-sgRNA:Design 1, Addgene # 159538). Oligonucleotides with spacer sequences for target genes were inserted into the sgRNA cassette by ligation into the BspQI and BsmBI cloning sites.

### Cloning of rAAV:HDR plasmids

Donor DNA fragments with target sites (AAV:HDR:cleaved) or without target sites (rAAV:HDR:uncleaved) were synthesized as gblocks (IDT) and cloned into the minimized Nme2Cas9+sgRNA AAV backbone plasmid following digestion of the latter with SalI restriction enzyme. The rAAV:HDR:uncleaved plasmids were cloned in NEB 5-alpha Competent *E. coli* cells (Catalog # C2987H), while rAAV:HDR:cleaved plasmids were cloned into the DH5α-AcrIIC4_Hpa_ *E. coli* strain that we constructed. Oligonucleotides with spacer sequences for target genes were inserted into the sgRNA cassette by ligation into the BspQI cloning sites.

### Creating *E. coli* DH5 cells stably expressing anti-CRISPR protein DH5α-AcrIIC4_*Hpa*_

Cells were created by λRed-promoted PCR-mediated recombination as previously described^78^. Briefly, *acrIIc4* _*Hpa*_, its promoter and Kan^R^ cassette were cloned from plasmid pUC57mini-AcrIIC4_*Hpa*_ (generously provided by Dr. Alan Davidson’s lab at the University of Toronto) into the pKIKOarsBKm (Addgene #46766) plasmid by restriction digestion and ligation into the *arsB* locus of DH5α. Cells harboring plasmid pKM208 (Addgene #13077) were cultured at 30°C to an O.D. of 0.35, induced with 1 mM IPTG, and harvested at an O.D. of 0.80. Cells were prepared for electroporation (washing with 10% glycerol) and transformed with 500 ng of gel-purified PCR product (*acrIIc4*_*Hpa*_, its promoter and Kan^R^). After incubation at 37°C overnight in kan-LB media, colonies were screened by PCR for the *acr* gene and the expected junction of the recombineered product with the *arsB* gene. Correct clones were passaged at 42°C to eliminate the pKM208 plasmid.

The rAAV:HDR:cleaved plasmids were linearized with BspQI restriction enzyme and spacer sequences with compatible overhangs were ligated into 100 ng of plasmid backbone using ElectroLigase® (NEB M0369) following the manufacturer’s protocol. For electro-transformation, 2 μL of ligation reaction was added to 40 μL of DH5α-AcrIIC4_*Hpa*_ *E. coli* and electroporated at 2500 V, 200 Ω, 25 μF, in a 2 mm gap cuvette using Gene Pulser Xcell Electroporation Systems [Bio-Rad #1652660].

### Vector production

Packaging of AAV vectors was done at the Viral Vector Core of the Horae Gene Therapy Center at the University of Massachusetts Medical School as described previously^64^. Constructs for HT-I studies were packaged in AAV8 capsids, whereas constructs for MPS-I studies were packaged in AAV9 capsids. A plasmid expressing anti-CRISPR protein (pEJS581; pCSDest2-AcrIIC4_*Hpa*_-FLAG-NLS; Addgene # 113436) was included in the triple-transfection packaging process to maintain intact rAAV:HDR:cleaved plasmids during production. Vector titers were determined by droplet digital PCR, and by gel electrophoresis followed by silver staining.

### Cell culture

HEK293T cells harboring Traffic Light Reporter Multi-Cas Variant 1 (TLR-MCV1)^81^, as well as Neuro2a cells (ATCC CCL-131), were cultured in Dulbecco’s Modified Eagle Media (DMEM, Thermo Fisher Scientific, Cat. No. 11965084); while MOLT-3 cells were cultured in RPMI-1640 medium. Both media were supplemented with 10% FBS (Thermo Fisher Scientific, Cat. No. 10438034) and 1% Penicillin/Streptomycin (GIBCO). Cells were incubated at a 37°C incubator with 5% CO_2_.

### Plasmid transfection

HEK293T cells and TLR-MCV1 cells were transfected using PolyFect Transfection Reagent (Qiagen, Cat No. 301105) with 400 ng of plasmid in a 24-well plate according to the manufacturer’s protocol. Neuro2a cells were transfected with 1000 ng of plasmid using Lipofectamine 3000 Reagent (Thermo Fisher Scientific, Cat. No. L3000015) following the manufacturer’s protocol. FACS analysis for the TLR-MCV1 cells were performed 3 days post-transfection using a Miltenyi MACSQuant VYB flow cytometer. HEK293T and Neuro2a cells were harvested 3 days post-transfection and genomic DNA was extracted using a DNeasy Blood and Tissue kit (Qiagen) according to the manufacturer’s protocol.

### Ribonucleoprotein nucleofection

Nme2Cas9 protein was expressed in Rosetta (DE3) cells and purified using a Ni^2+^-NTA agarose column (QIAGEN) as previously described^24^. Wild-type (wt) gRNA was in vitro transcribed using the HiScribe T7 High Yield RNA Synthesis Kit (New England Biolabs) according to the manufacturer’s protocol. Truncated guide (Nme.sgRNA-100) was chemically modified with 2’-*O*-methyl analogs and 3’-phosphorothioates in 5’ and 3’ terminal RNA residues, and was synthesized by Synthego (Redwood City, CA, USA). RNP complex was electroporated into HEK293T or MOLT-3 cells using the Neon transfection system. Briefly, 40 picomoles (HEK293T and MOLT-3 cells) of 3xNLS-Nme2Cas9 and 50 picomoles (HEK293T and MOLT-3 cells) of T7-transcribed or chemically synthesized sgRNA was assembled in buffer R along with 100,000 HEK293T cells or 200,000 MOLT-3 cells and electroporated using 10 μL tips. Electroporation parameters (voltage, width, number of pulses) were 1150 V, 20 ms, 2 pulses for HEK293T cells; 1600 V, 10 ms, 3 pulses for MOLT-3 cells.

### TIDE analysis, amplicon sequencing and indel analysis

PCR was carried out with TIDE or amplicon sequencing primers as shown in the Supplementary Table using High Fidelity 2x PCR Master Mix (New England Biolabs). PCR products were purified using the DNA Clean & Concentrator-100 (Zymo, Cat. No. D4029) and sent for Sanger sequencing using a TIDE forward primer (Supplementary Table). Indel readouts were obtained using the TIDE web tool (https://tide-calculator.nki.nl/) as previously described^27^. For amplicon sequencing, PCR was carried out using amplicon sequencing primers shown in the Supplementary Table. Equimolar amounts of DNA were amplified with a universal forward primer and an indexed reverse primer (98 °C, 15 s; 61 °C, 25 s; 72 °C, 18 s; nine cycles) to ligate the TruSeq adaptors. The resultant amplicons were purified using Ampure beads. Libraries were sequenced for 600 cycles (300/300 PE) on a MiSeq (*Fah* locus), and for 300 cycles (SE) on a MiniSeq (*Idua* locus).

FASTQ files were analyzed using CRISPResso2 with parameters specifying guide, amplicon and expected HDR sequences. A no-guide sample was used as the negative control to determine background. AAV integration was quantified as previously described^87^ using BWA-MEM and *samtools* to align sequenced amplicons files to the *Idua* locus and AAV vector sequence.

### SMRT sequencing

Amplicon libraries were constructed using locus-specific primers as shown in the Supplementary Table using Q5® High-Fidelity DNA Polymerase (New England Biolabs). One primer of each pair contains an 8-nucleotide UMI and both contain adapter overhangs to multiplex the library using Bar. Univ. F/R Primers Plate-96v2 (Pacbio part no. 101-629-100).

Forward and reverse primers were 1.1 kb and 1.9 kb from gRNA-I and gRNA-II Cas9 DSB sites, respectively. Each sample was amplified by linear amplification (98°C for 40 s, followed by 10 cycles of 98 °C for 30 s, 64 °C for 20 s and 72°C for 75 s) using the forward primers. A second PCR amplification was performed using a universal PacBio primer and a gene-specific reverse primer at 98°C for 30 s, followed by 30 cycles of 98°C for 15 s, 64°C for 20 s and 72°C for 75 s, and extended at 72°C for 5 m. PCR products were diluted (1:100) and amplified using PacBio barcoded primers at 98°C for 30 s, followed by 30 cycles of 98°C for 15 s, 64°C for 20 s and 72°C for 75 s, and extended at 72°C for 12 m. Equimolar amounts of the products were combined together and purified by AMPure PB beads (PacBio, part no. 100-265-900) following the manufacturer’s recommendation. The libraries were sequenced on an Pacific Biosciences Sequel II Instrument in a 10-hour collection SMRTCell™. Circular consensus sequence (ccs) reads generated by SMRT Link v8 using default parameters in fastq format were processed and analyzed using the Galaxy web platform at usegalaxy.org^88^ with custom workflows, unless specified. Briefly, reads were demultiplexed using *FASTQ/A Barcode Splitter*, and converted to fasta to calculate and count read lengths within each library (Supplementary Fig. 2). To assign reads that represent segmental deletions, inversions, or indel events, reads were first re-IDed by appending each sequence identifier with the UMIs using fastp with (-U). Duplicated UMIs were removed from the analysis. Reads with mapping qualities >20 were then uniquely aligned using BWA-MEM to reference sequences representing the wild-type loci and the predicted sequences following segmental deletion or inversion. The counts for reads aligning to the specified references were displayed as a percentage of all mapped reads. To categorize reads that mapped to the wild-type reference as either bearing indel events or unedited genomes at indicated target loci, reads were trimmed to ±100 nt surrounding each target PAM by *Cutadapt*. Trimmed reads were then converted to fasta and clustered by *UCLUST*^*89*^, a centroid-based clustering algorithm. Singlets were discarded and treated as reads bearing sequencing errors. Clustered reads using size reporting (-sizeout) were categorized into unedited and edited groups at each target site or both sites for dual-guide vectors and displayed as a percentage of all reads mapping to the respective wild-type sequence. To assess target loci that harbor vector genome integration events, fastq reads were mapped to the relevant vector genome reference and expressed as a percentage of all reads.

### RNA sequencing

Total RNA was isolated from liver tissues using TRIzol reagent and separated by chloroform. RNA-seq library preparation was performed with 1ug of total RNA per sample using the TruSeq Stranded mRNA Library prep kit (Illumina, 20020595) following the manufacturer’s protocol. Libraries were made for the following mice: 3 PBS treated *Fah*^*neo/neo*^, 8 dual-sgRNA Design 1 treated *Fah*^*neo/neo*^, 8 dual-sgRNA Design 4 treated *Fah*^*neo/neo*^, 3 healthy C57BL/6, 4 dual-sgRNA Design 1 treated C57BL/6, 4 dual-sgRNA Design 4 treated C57BL/6, 3 PBS treated *Fah*^*PM/PM*^, 3 rAAV:HDR:FahDonor:ncSpacer treated *Fah*^*PM/PM*^, 3 rAAV:HDR:ncDonor:FahSpacer treated *Fah*^*PM/PM*^, 10 rAAV:HDR:uncleaved treated *Fah*^*PM/PM*^, and 10 rAAV:HDR:cleaved treated *Fah*^*PM/PM*^. Libraries were sequenced on an Illumina NextSeq 500 with single-end 75 nucleotide reads and an average of 19.88 million reads. All downstream analyses were performed with the Ensembl GRCm38.p6 (GCA_000001635.8) *Mus musculus* genome adapted to include the Nme2Cas9 mRNA sequence. Transcripts Per Million (TPM) were calculated with alignment-free quantification using Kallisto v0.45.0. Reads were mapped to the mm10 genome using STAR v2.7.5a in chimeric mode to identify chimeric junction reads and HTSeq v.0.10.0 was used to quantify read counts for each gene. Differential expression analyses were performed with DESeq2 release 3.11 after filtering out genes with less than 10 read counts across all samples in the comparison. Gene ontology analyses were conducted with a custom script to test for enrichment of functional annotations among significantly differentially expressed genes, in order to avoid significant gene ontology terms with overlapping gene sets. Specifically, the script uses gene ontology annotation databases from the Gene Ontology Consortium to perform iterative enrichment analyses and *p* values are computed using a Fisher-exact test and then corrected using an Benjamini-Hochberg multiple test correction.

### HPD western blotting

Liver tissue fractions were ground, resuspended in 200 μL of RIPA lysis buffer and allowed to lyse for 15 m on ice. Measurements of total protein content was determined by Pierce™ BCA Protein Assay Kit (Thermo-Scientific) following the manufacturer’s protocol. A total of 25 μg of protein was loaded onto a 4–20% Mini-PROTEAN® TGX™ Precast Gel (Bio-Rad) and transferred onto a PVDF membrane. After blocking with 5% Blocking-Grade Blocker solution (Bio-Rad) for 2 h at room temperature, membranes were incubated with rabbit anti-HPD (Sigma _Hpa_038322, 1:600) overnight at 4 °C. Membranes were washed three times in TBST and incubated with horseradish peroxidase (HRP)-conjugated goat anti-rabbit (Bio-Rad 1,706,515, 1:10,000) secondary antibodies for 2 h at room temperature. The membranes were washed four times in TBST and visualized with Clarity™ western ECL substrate (Bio-Rad) using a Bio-Rad ChemiDoc MP imaging system. Membranes were stripped using Restore™ Western Blot Stripping Buffer (Cat. number: 21059) following the manufacturer’s protocol and blocked with 5% Blocking-Grade Blocker solution (Bio-Rad) for 2 h at room temperature. The membranes were incubated with rabbit anti-GAPDH (Abcam ab9485, 1:2,000) overnight at 4°C, washed three times in TBST, and incubated with HRP-conjugated goat anti-rabbit (1:4,000) secondary antibodies for 2 h at room temperature. The membranes were washed four times in TBST and visualized with Clarity™ western ECL substrate (Bio-Rad) as above.

### Histological analysis

Liver tissues were harvested, and sections were placed in biopsy cassettes and fixed in 10% buffered formalin overnight before changing into 70% ethanol solution. FAH immunohistochemistry (IHC) was performed using anti-FAH antibody (Abcam, Cat. No. ab81087; dilution 1:400), whereas HPD IHC was performed using anti-HPD antibody (Sigma _Hpa_038322; dilution 1:30). IHC and hematoxylin and eosin (H&E) staining were performed by the Morphology Core at University of Massachusetts Medical School following standard procedures.

### Reverse transcription PCR

Total RNA was isolated from liver tissues using TRIzol reagent and separated by chloroform. Reverse transcription reactions were carried out using SuperScript™ III First-Strand Synthesis System (Thermo Fisher Scientific; 18080051) following the manufacturer’s protocol. The complementary DNA was used to quantify *Fah* gene expression in a qPCR reaction using primers shown in the Supplementary Table using PowerUp SYBR Green Master Mix (Applied Biosystems, Cat. No. A25742). All reactions were normalized to *gapdh* and relative quantification in gene expression was determined using the 2^-ΔΔCt^ method. PCR products were resolved by agarose gel electrophoresis.

### Quantitative PCR for rAAV copy number

Genomic DNA from mouse tissue was used to quantify rAAV copy number using primers shown in the Supplementary Table using PowerUp SYBR Green Master Mix (Applied Biosystems, Cat. No. A25742). All reactions were normalized to *gapdh* control and relative quantification of rAAV copies was determined using the 2^-ΔΔCt^ method relative to PBS-injected cohorts.

### Serum aspartate transaminase (AST) and alanine transaminase (ALT) assays

Blood was collected from mice by cardiac puncture immediately before euthanasia, and sera were isolated using a serum separator (BD, Cat. No. 365967) and stored at −80°C. AST and ALT levels were determined using Cobas Clinical Chemistry Analyzer at the MMPC-University of Massachusetts Medical School Analytical Core (RRID:SCR_015365). Two mice in the PBS- and rAAV:ncDonor:FahSpacer negative control HT-I cohorts were hemolyzed, so we were able to measure ALT/AST for only one mouse. In the rest of the groups, data are presented as mean values ± s.e.m. (n = 3-7 mice per group).

### Iduronidase activity assay

The iduronidase activity assay was performed as described previously^86,90,91^ with minor modifications. Tissues were homogenized in ice-cold T-PER protein extraction reagent (Thermo Fisher Scientific, Cat. No. 78510) with protease inhibitor (Roche, Cat. No. 4693159001) using TissueLyser II (Qiagen). Quantification of total protein concentration was assessed by the bicinchoninic acid (BCA) method (Pierce, Cat. No. 23225). No more than 80 μg of total protein was used in enzymatic reactions (100 μL of total reaction volume), which includes sodium formate buffer, pH 3.5 (130 mM), D-saccharic acid 1,4-lactone monohydrate (0.42 mg/mL, Sigma-Aldrich, Cat. No. S0375), and 4MU-iduronic acid (0.12 mM, Santa Cruz Biotechnology, Cat. No. sc-220961). The reaction was incubated under 37°C for 24-48 h, and quenched with glycine buffer (pH 10.8). The fluorescence of released 4MU (excitation wavelength: 365 nm; emission wavelength: 450 nm) was detected using a fluorescence plate reader (BioTek) and compared against a standard curve generated using 4MU (Sigma-Aldrich, Cat. No. M1381). The iduronidase specific activity was calculated as 4MU released (pmole) per milligram of total protein per hour.

### Glycosaminoglycan (GAG) assay

The glycosaminoglycan (GAG) assay was performed using a protocol previously described^91^. Tissues were homogenized in a mixture of chloroform and methanol (2:1) using TissueLyser II (Qiagen) and dried in a Vacufuge (Eppendorf) to remove fat. The dried and defatted tissue was weighed and digested using papain (Sigma-Aldrich, Cat. No. P3125) at 60°C overnight. The supernatant was used in the Blyscan assay to quantify GAG content using chondroitin-4-sulfate as standard (Accurate Chemical, Cat. No. CLRB1000). Levels of GAG were calculated as GAGs (microgram) per milligram of dried, defatted tissue.

### Humoral α-Nme2Cas9 immune response

Nme2Cas9-specific humoral immune response was measured as previously described^27^. Briefly, a 96-well plate (Corning) was coated with 0.5 μg of recombinant Nme2Cas9 protein suspended in 1× coating buffer (Bethyl E107). The plate was incubated for 12 h at 4°C with shaking, then washed three times using 1x Wash Buffer (Bethyl E106). The plate was blocked with blocking buffer (Bethyl E104) for 2 h at room temperature. After washing three times, diluted sera (1:40 in dilution buffer consisting of 1L of TBS, 10 g of BSA and 5 mL of 10% Tween 20) from mice injected with rAAV:Nme2Cas9 were added to each well in duplicates and incubated at 4°C for 6 h. The plate was washed three times, and goat anti-mouse HRP-conjugated antibody (100 μL; 1:100,000 dilution; Bethyl A90-105P) was added to each well. The plates were incubated at room temperature for 1 h, then washed four times. To detect immune response, the plate was developed by adding 100 μL of TMB One Component HRP Substrate (Bethyl E102). After incubating in the dark for 30 m at room temperature, Stop Solution (Bethyl E115; 100 μL per well) was directly added. Absorbance was measured at 450 nm using a BioTek Synergy HT microplate reader.

## Statistical analysis

All data are presented as mean values ± s.e.m. *p* values are calculated using Student’s t-test (2-sided) and one-way ANOVA using Grap_Hpa_d Prism.

### Reporting Summary

Further information on research design is available in the Nature Research Reporting Summary linked to this article.

## Data availability

Sequencing data that support the findings of this study have been deposited in SRA under project PRJNA667456.

## Code availability

All analysis was performed using publicly available programs and the parameters indicated in the Methods section.

## Acknowledgments

We thank members of the Sontheimer, Gao, Wang, Xue and Wolfe labs for helpful advice and discussions, and Alan Davidson for anti-CRISPR plasmid. We are grateful to the UMass Medical School Mouse Phenotyping Center, Viral Vector Core, Sequencing Core, Morphology Core and Animal Medicine staff for their expertise and assistance. We also thank Krishna Ghanta for his assistance with SMRT sequencing library design, Jordan Smith for help with animal handling, and Jun Xie for AAV packaging expertise. Multiple figures of this manuscript were created with BioRender.com. Support for this work was provided by the National Institute of Diabetes and Digestive and Kidney Diseases of the National Institutes of Health (F31DK120333 to R.I.), as well as NIH grants R01NS076991 and U19AI149646 to G.G., P01HL131471 and UG3HL147367 to G.G. and W.X., DP2HL137167 to W.X., R35GM133762 to A.A.P., R01GM125797 to E.J.S., and R01GM115911 and UG3TR0228 to S.A.W. and E.J.S. Additional support was provided by funds from SLC61A Connect to A.A.P. and by grants from the American Cancer Society (129056-RSG-16-093) and the Cystic Fibrosis Foundation to W.X.

## Author Contributions

R.I. and E.J.S. conceived the study and designed experiments with the participation of all authors. R.I. engineered and tested plasmids and AAV vectors, processed mouse tissues, assembled sequencing libraries, and performed all Western blot, qRT-PCR, qPCR, tissue imaging, and immune response assays. A.M. designed and tested truncated sgRNAs, and A.M. and R.I. validated truncated sgRNAs in cellular editing. A.M. and T.R. performed computational analyses of short-read amplicon libraries. P.W.L.T. analyzed vector DNA and analyzed SMRT long-read sequencing data. E.M. and S.M. assisted in the analysis of editing outcomes. J.W. handled MPS-I mice and performed IDUA and GAG assays. N.J. and E.K. prepared RNA-Seq libraries, and N.J., E.K. and A.A.P. analyzed RNA-seq data. S.N. constructed *E. coli* strains expressing anti-CRISPR protein. Y.C. and E.T. assisted with animal experiments. S.A.W., D.W., A.A.P., W.X., G.G. and E.J.S. oversaw experimental design and interpretation and all authors assisted in data interpretation. R.I. and E.J.S. wrote the manuscript and all authors edited the manuscript.

---

**Supplementary Fig. 1.**
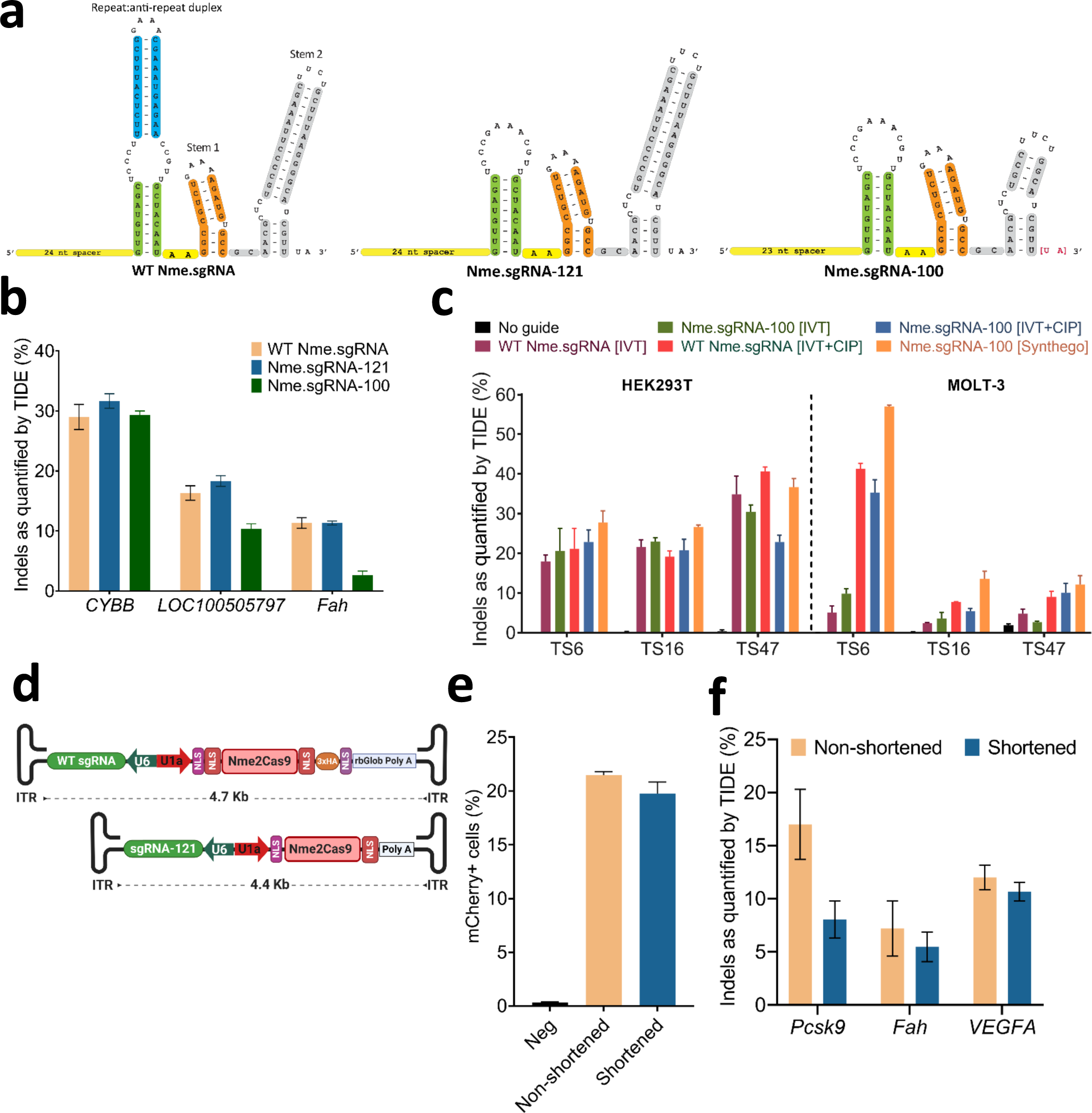
sgRNA truncation and AAV:Nme2Cas9 vector minimization. **a**, Illustration of the 145-nt, full-length Nme.sgRNA, the truncated 121-nt Nme.sgRNA-121 and the further truncated 100-nt Nme.sgRNA-100. **b**, Comparison of editing efficiency with WT Nme.sgRNA, Nme.sgRNA-121 and Nme.sgRNA-100 by plasmid transfection at *CYBB, LOC100505797* (in HEK293T) and *Fah* (in Neuro2a) genomic loci as estimated by TIDE analysis. **c**, Comparison of RNP editing efficiencies using full-length Nme.sgRNA and Nme.sgRNA-100, produced as T7 RNA polymerase transcripts (*in vitro* transcribed, IVT) with and without treatment with phosphatase (CIP), along with chemically synthesized, commercial Nme.sgRNA-100 guides. Nme2Cas9 RNPs targeting three genomic sites (TS6, TS16 and TS47) were electroporated into HEK293T and T lymphoblast MOLT-3 cells, and editing was assessed by TIDE analysis. **d**, Schematic of the ∼4.7 kb (top) and the minimized ∼4.4 kb (bottom) AAV vectors expressing Nme2Cas9 and a 121nt sgRNA. ITR, inverted terminal repeats. Nme2Cas9 includes SV40 and nucleoplasmin NLSs on the N and C termini, respectively. **e**, Comparison of editing efficiencies of the non-shortened 4.7 Kb and the shortened 4.4 Kb AAV:Nme2Cas9 constructs by plasmid transfection in TLR-Multi-Cas-Variant 1 (MCV1) lentivector-transduced HEK293T cells, as measured by flow cytometry. **f**, Comparison of editing efficiencies of the ∼4.7 Kb and the shortened ∼4.4 Kb AAV:Nme2Cas9 plasmids tested by transfection in Neuro2a cells (*Pcsk9* and *Fah*) and HEK293T (*VEGFA*) and estimated by TIDE analysis. All data are presented as mean values ± s.e.m. (n = 3 biological replicates).

**Supplementary Fig. 2.**
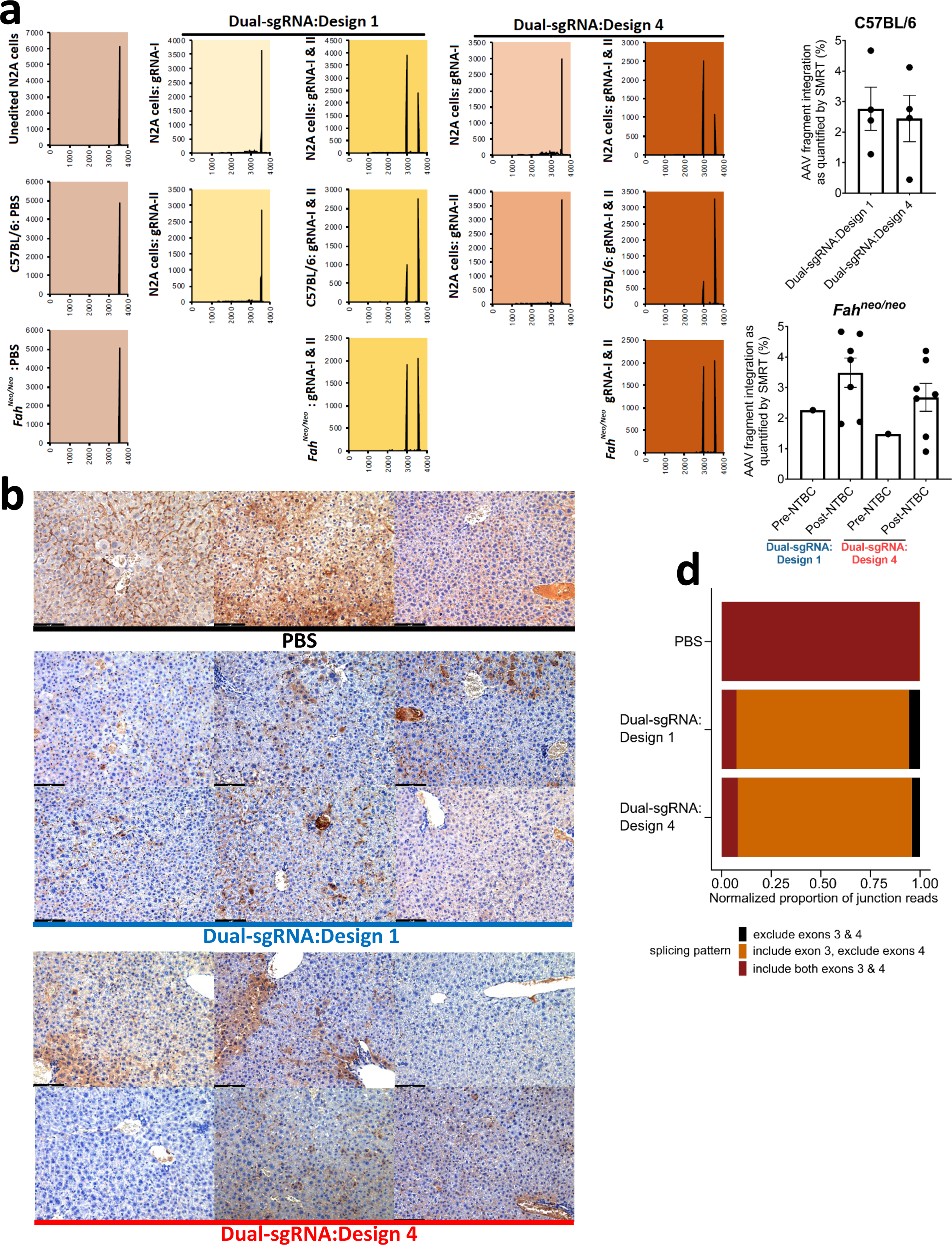
*In vivo* editing using dual-sgRNA rAAV:Nme2Cas9 vectors. **a**, Representative length distribution plots for mapped SMRT reads indicating the presence of segmental deletions in Neuro2a (N2A) cells after plasmid transfection, or in the livers of C57BL/6 and *Fah*^*neo/neo*^ mice after AAV8 delivery. **b**, Anti-HPD immunostaining in liver tissues from *Fah*^*neo/neo*^ mice injected with PBS or with AAV8:Nme2Cas9 Dual-sgRNA:Designs 1 and 4 vectors. **c**, rAAV fragment integration as detected by SMRT sequencing analysis. Data are presented as mean values ± s.e.m. (n = 4 in C57BL/6 cohort; n = 1 in pre-NTBC *Fah*^*neo/neo*^ cohort; n = 7 in post-NTBC *Fah*^*neo/neo*^ cohort). **d**, Proportions of exon-exon junction reads from RNA-seq data that support the inclusion of both exons 3 and 4 (red), inclusion of only exon 3 (orange), and exclusion of both exons 3 and 4 (black) of *Hpd* in PBS-, Dual-sgRNA:Design 1-, and Dual-sgRNA:Design 4-treated *Fah*^*neo/neo*^ mice.

**Supplementary Fig. 3.**
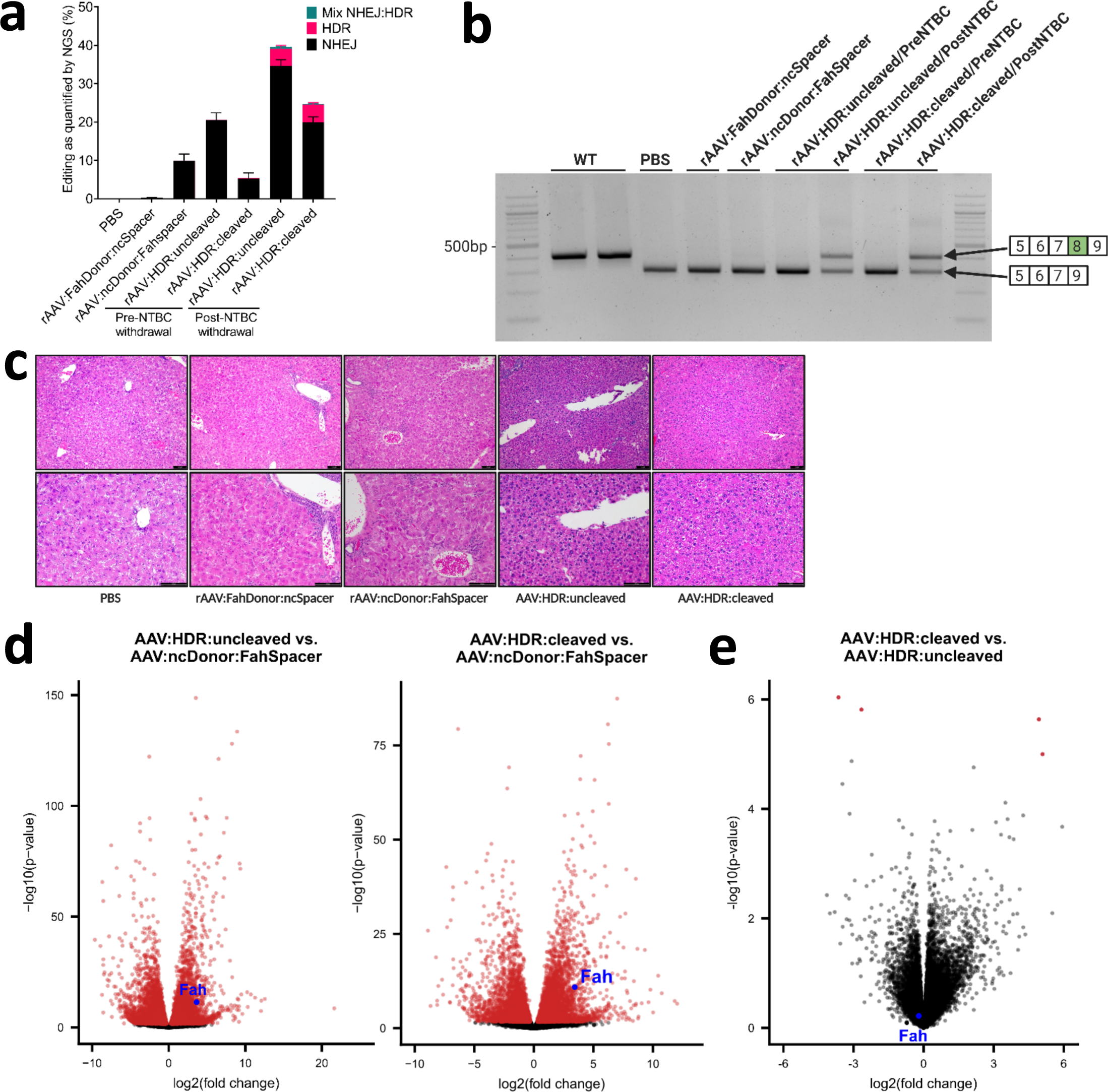
Improved molecular and pathological phenotypes of HT-I mice after treatment with rAAV:HDR vectors. **a**, Stacked histograms showing the percentages of NHEJ, HDR and imprecise NHEJ:HDR mix at the *Fah* editing site in livers of HT-I mice pre- and post-NTBC withdrawal, as measured by NGS sequencing of PCR amplicons. **b**, Agarose gel image showing the detection of RT-PCR products of *Fah* mRNA in liver lysate. The lower band (∼300 bp) is from products with exon 8 skipped, while the ∼400 bp band is from products that include exon 8. **c**, Representative H&E staining of liver from HT-I mice injected with PBS or negative control (rAAV:FahDonor:ncSpacer and rAAV:ncDonor:FahSpacer) vectors, or with rAAV:HDR:cleaved and -uncleaved vectors. Scale bars are 100 mm for the upper panels and 20 mm for the lower panels. **d**, Volcano plots showing significantly differentially-expressed genes (red, FDR <= 5%) between AAV:HDR:uncleaved and AAV:ncDonor:FahSpacer *Fah*^*PM/PM*^ mice (left) and AAV:HDR:cleaved and AAV:ncDonor:FahSpacer *Fah*^*PM/PM*^ mice (right). **e**, Volcano plot showing the paucity of significantly differentially expressed genes (red, FDR <= 5%) between AAV:HDR:uncleaved and AAV:HDR:cleaved *Fah*^*PM/PM*^ mice. Data are presented as mean values ± s.e.m. (n = 3 in PBS, rAAV:FahDonor:ncSpacer, rAAV:ncDonor:FahSpacer and pre-NTBC withdrawal AAV.HDR:cleaved and - uncleaved cohorts; n = 7 in post-NTBC withdrawal rAAV:HDR:cleaved and -uncleaved cohorts).

**Supplementary Fig. 4.**
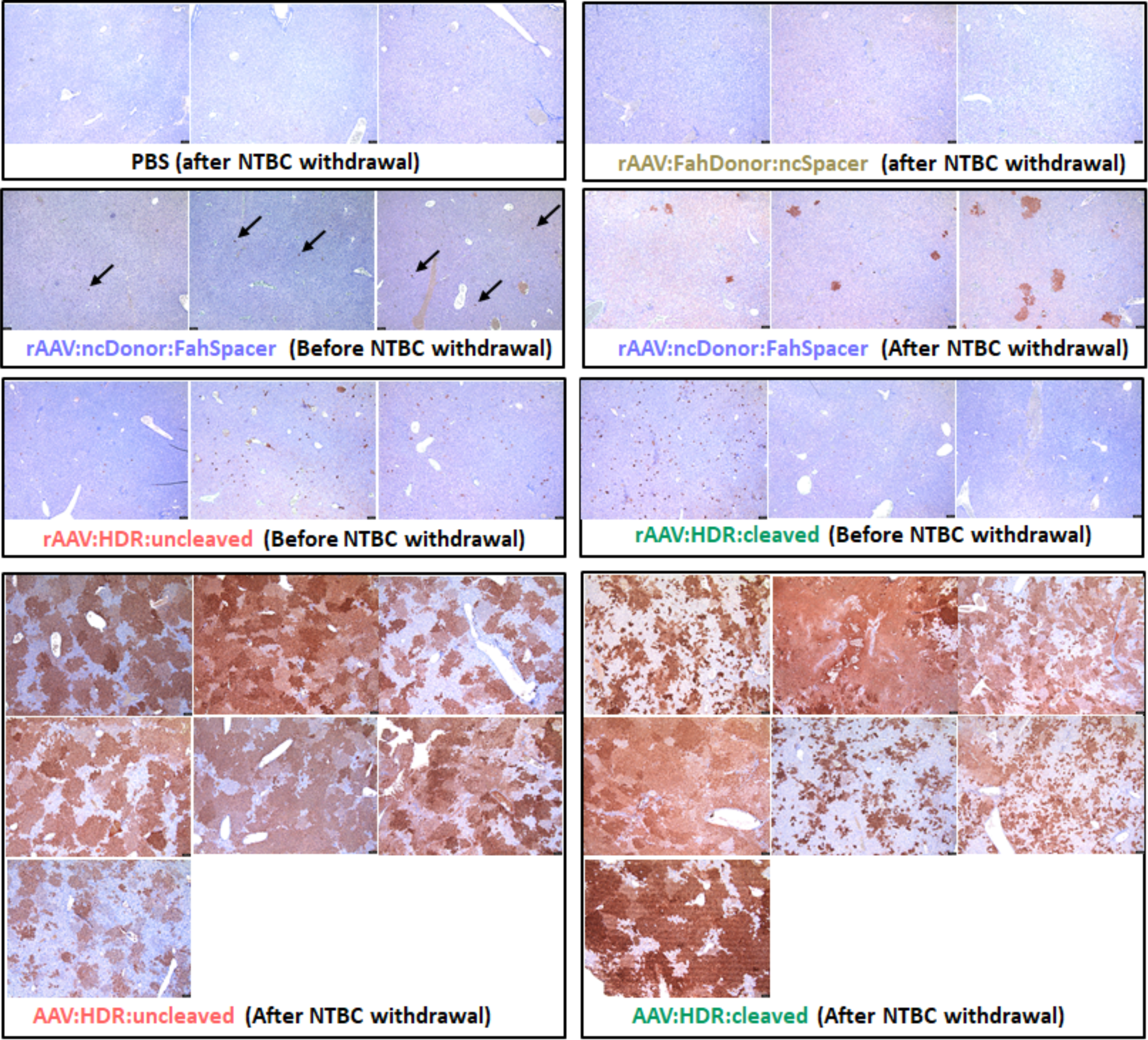
Anti-FAH immunostaining in liver tissues of all negative-control and treated cohorts. Vectors and drug regimens are indicated for each cohort. Scale bar, 100 mm.

**Supplementary Fig. 5.**
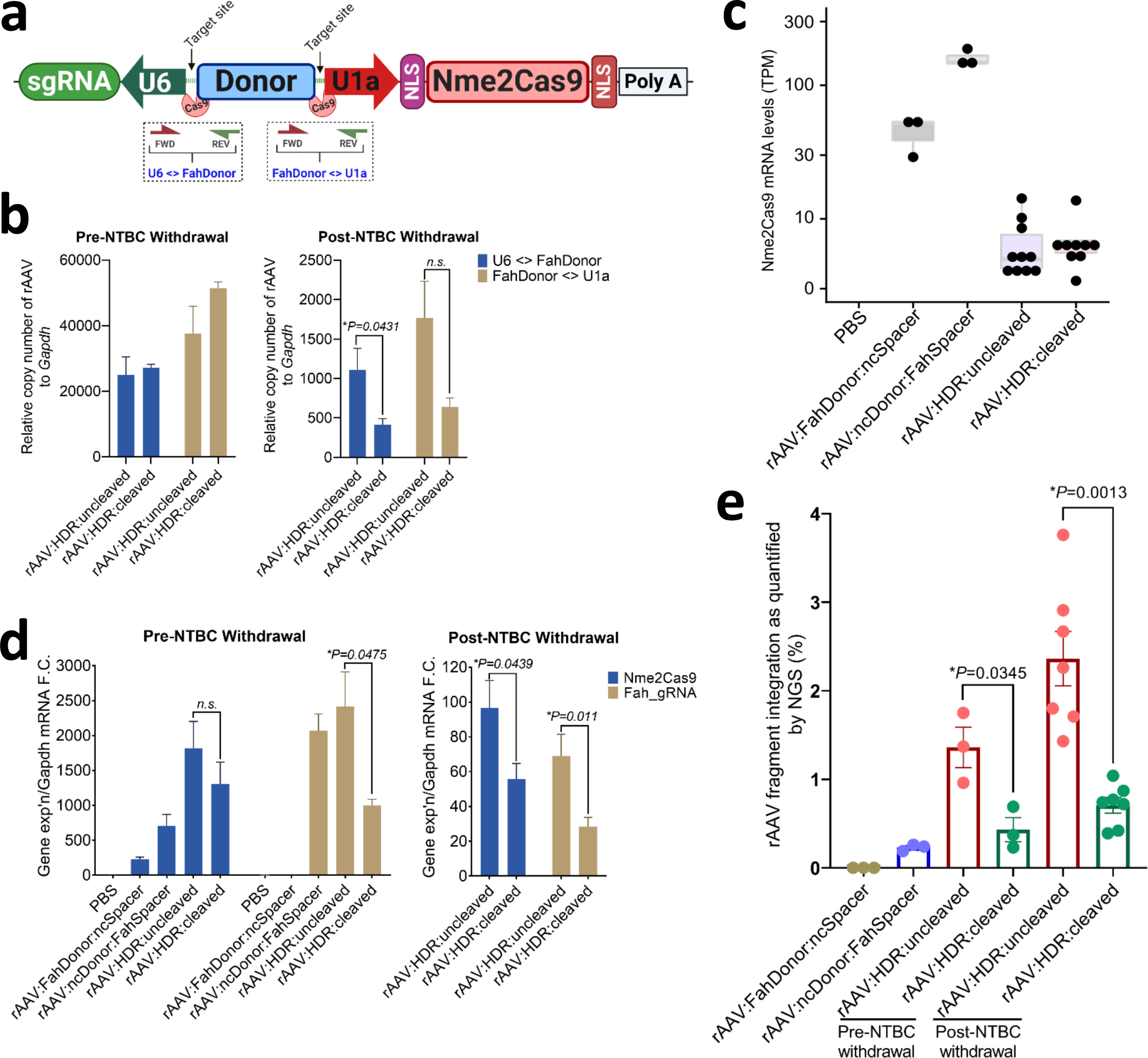
Reduced rAAV copy number, Nme2Cas9 mRNA expression, and sgRNA expression in HT-I mice treated with rAAV:HDR:cleaved vectors. **a**, Schematic of qPCR assays to quantitate rAAV genome segment copy number in HT-I mouse liver samples. Primer pairs [U6<>FahDonor] and [FahDonor<>U1a] are indicated underneath. **b**, rAAV copy numbers in HT-I mouse liver tissues. rAAV copies are significantly reduced in rAAV:HDR:cleaved cohorts after NTBC withdrawal using the [U6<>FahDonor] primer pair (left); the lower values observed with the [FahDonor<>U1a] primer pair do not reach statistical significance. **c**, Steady-state mRNA levels from RNA-seq (polyA-enriched, transcripts per million) of Nme2Cas9 mRNA in the liver of *Fah*^*PM/PM*^ mice. **d**, qRT-PCR analyses with total RNA showing reduced Nme2Cas9 mRNA and sgRNA expression in the rAAV:HDR:cleaved cohort after NTBC withdrawal. **e**, Reduced fragment integration of rAAV vectors in the *Fah* locus in cohorts injected with the rAAV:HDR:cleaved vector, as detected by NGS analysis. Data are presented as mean values ± s.e.m. (n = 3 in PBS, rAAV:FahDonor:ncSpacer, rAAV:ncDonor:FahSpacer and pre-NTBC withdrawal AAV:HDR:cleaved and -uncleaved cohorts; n = 7 in post-NTBC withdrawal rAAV:HDR:uncleaved and - uncleaved cohorts).

**Supplementary Fig. 6.**
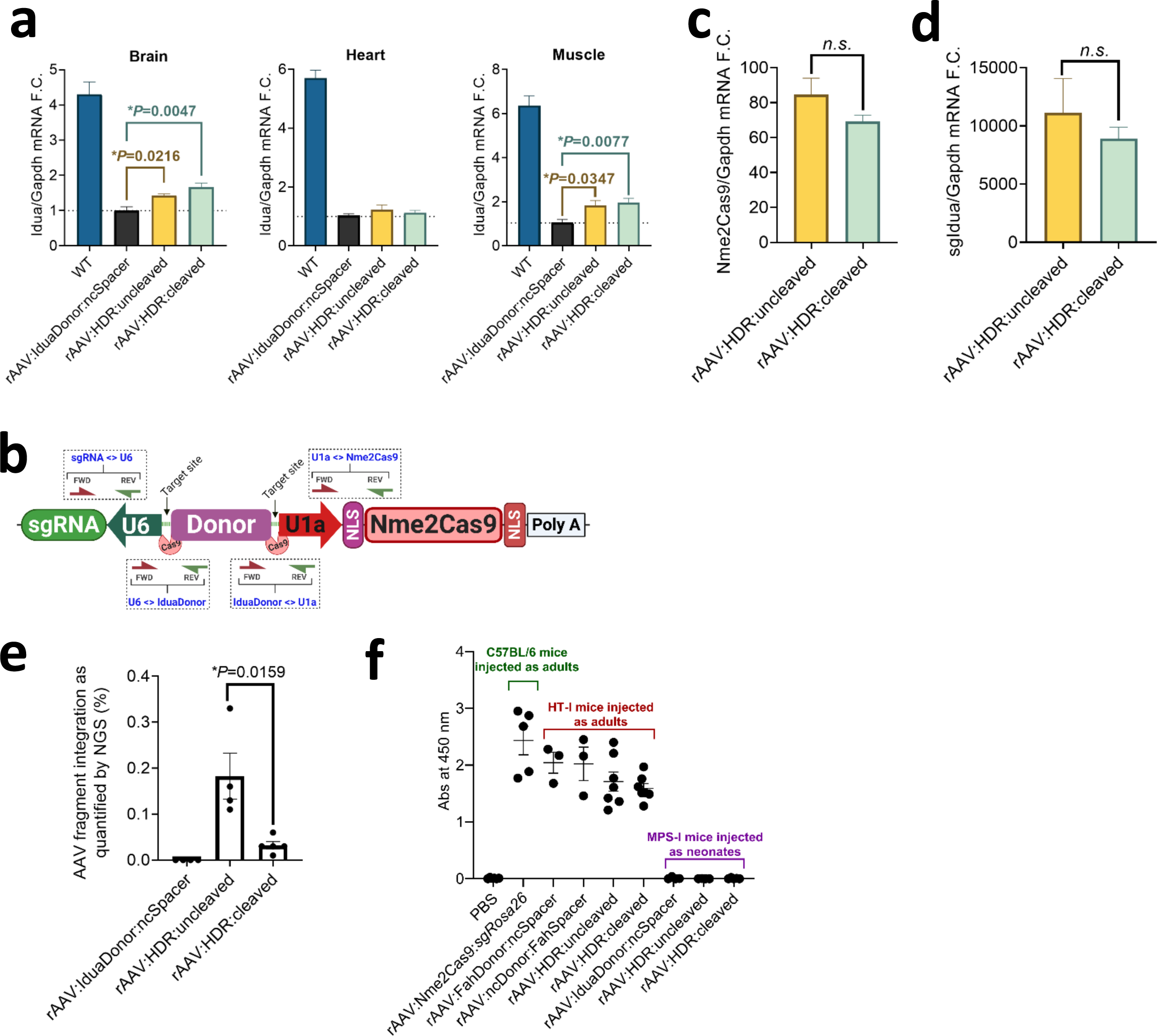
Reduced rAAV copy number, Nme2Cas9 mRNA expression, and sgRNA expression in MPS-I mice treated with rAAV:HDR:cleaved vectors. **a**, qRT-PCR data showing increase in *Idua* mRNA in the brain and muscle tissues as normalized to *Gapdh* mRNA. **b**, Schematic of qPCR assays to quantitate rAAV genome segment copy number in MPS-I mouse liver samples. Primer pairs [sgRNA<>U6], [U6<>IduaDonor], [IduaDonor<>U1a] and [U1a<>Nme2Cas9] are indicated underneath. **c**,**d** Animals treated with rAAV:HDR:cleaved vectors exhibit reduced Nme2Cas9 mRNA (**c**) and sgRNA (**d**) expression levels as measured by qRT-PCR, though reductions do not reach statistical significance. **e**, Reduced fragment integration of rAAV vectors in the *Idua* locus in livers of cohorts injected with the rAAV:HDR:cleaved vector, as detected by NGS analysis. **f**, Humoral IgG1 immune response to Nme2Cas9 *in vivo* is significantly reduced in rAAV-treated, neonate-injected mice compared to adult-injected cohorts. Data are presented as mean values ± s.e.m. (n = 3-5 biological replicates).

## Supplementary Note

Nucleotide sequence of all plasmids

### pEJS1089 mini-AAV.sgRNA.Nme2Cas9 construct

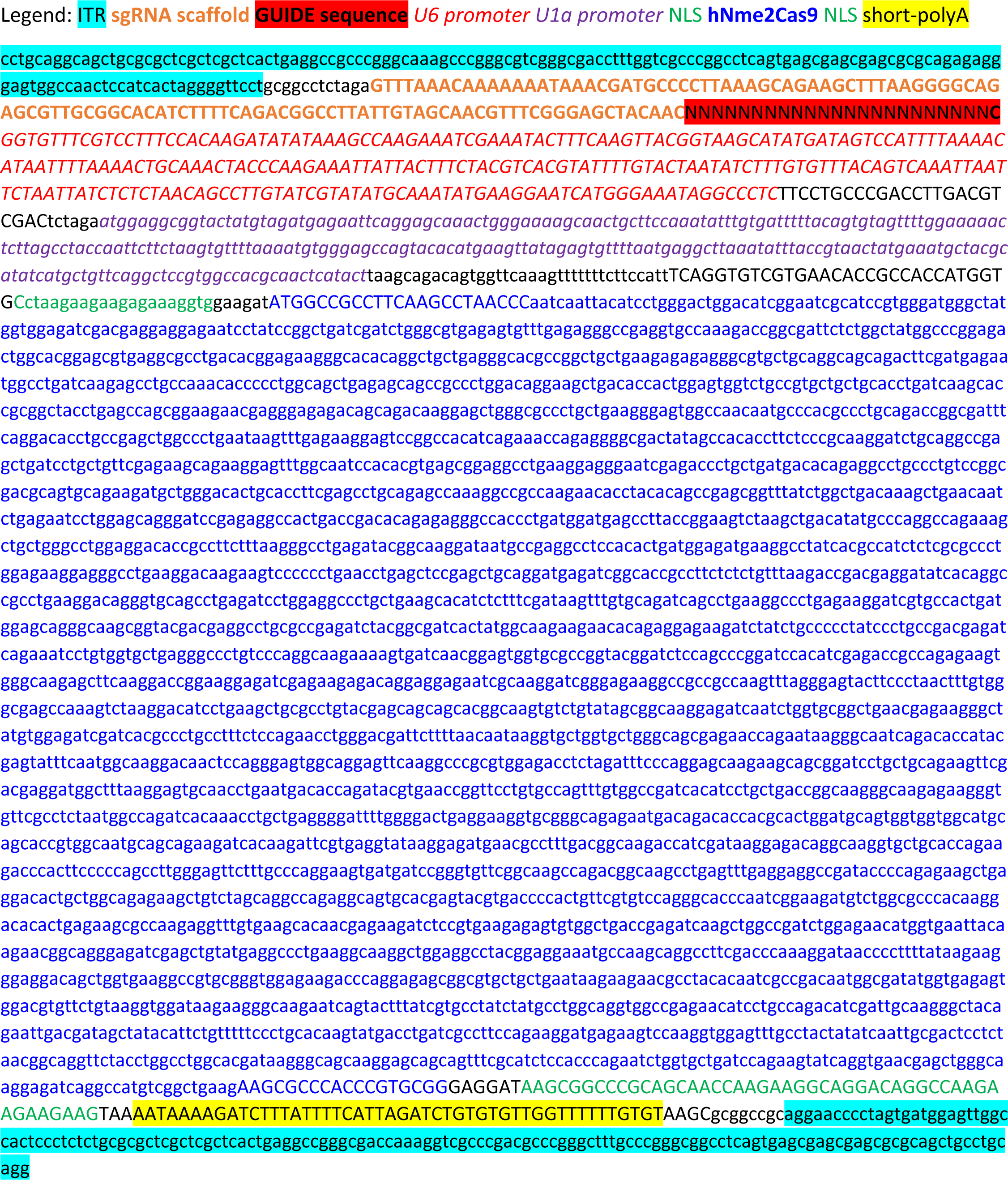

### pEJS1096: Dual-sgRNA.Design 1 construct

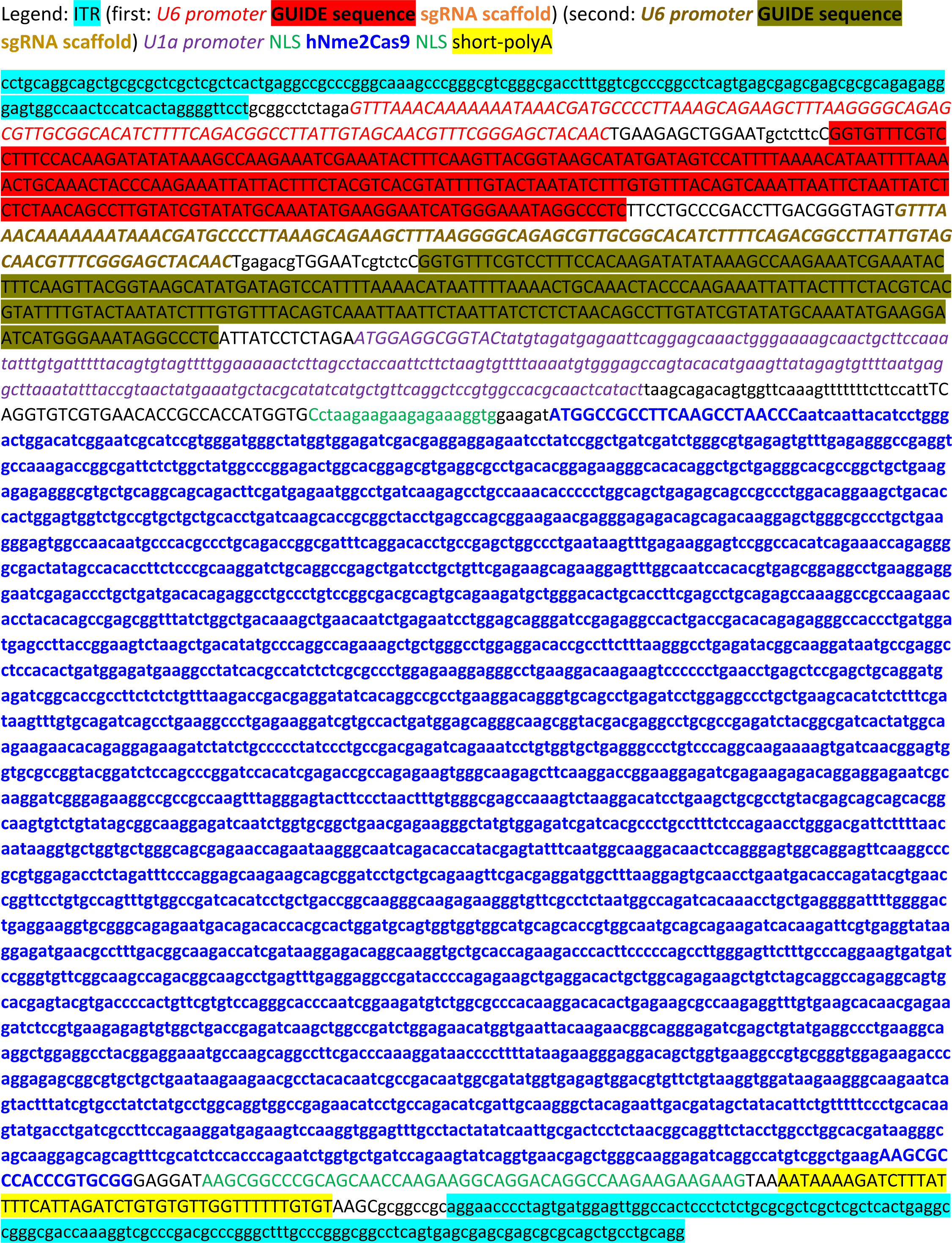

### pEJS1099: Dual-sgRNA.Design 4 construct

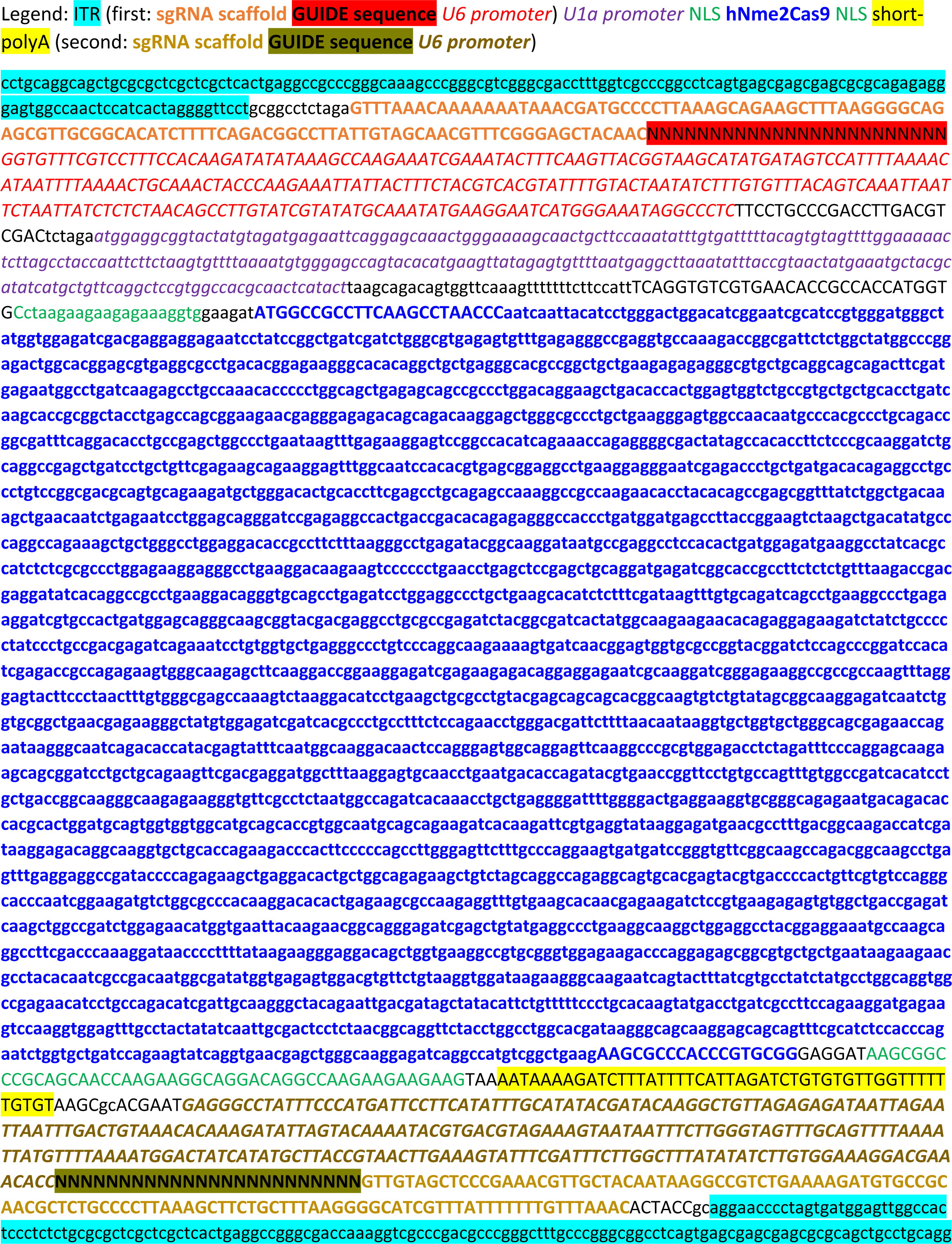

### AAV:HDR:uncleaved-Design B Plasmid to correct *eGfp* in HEK293T TLR-Multi-Cas-Variant 1 (MCV1)

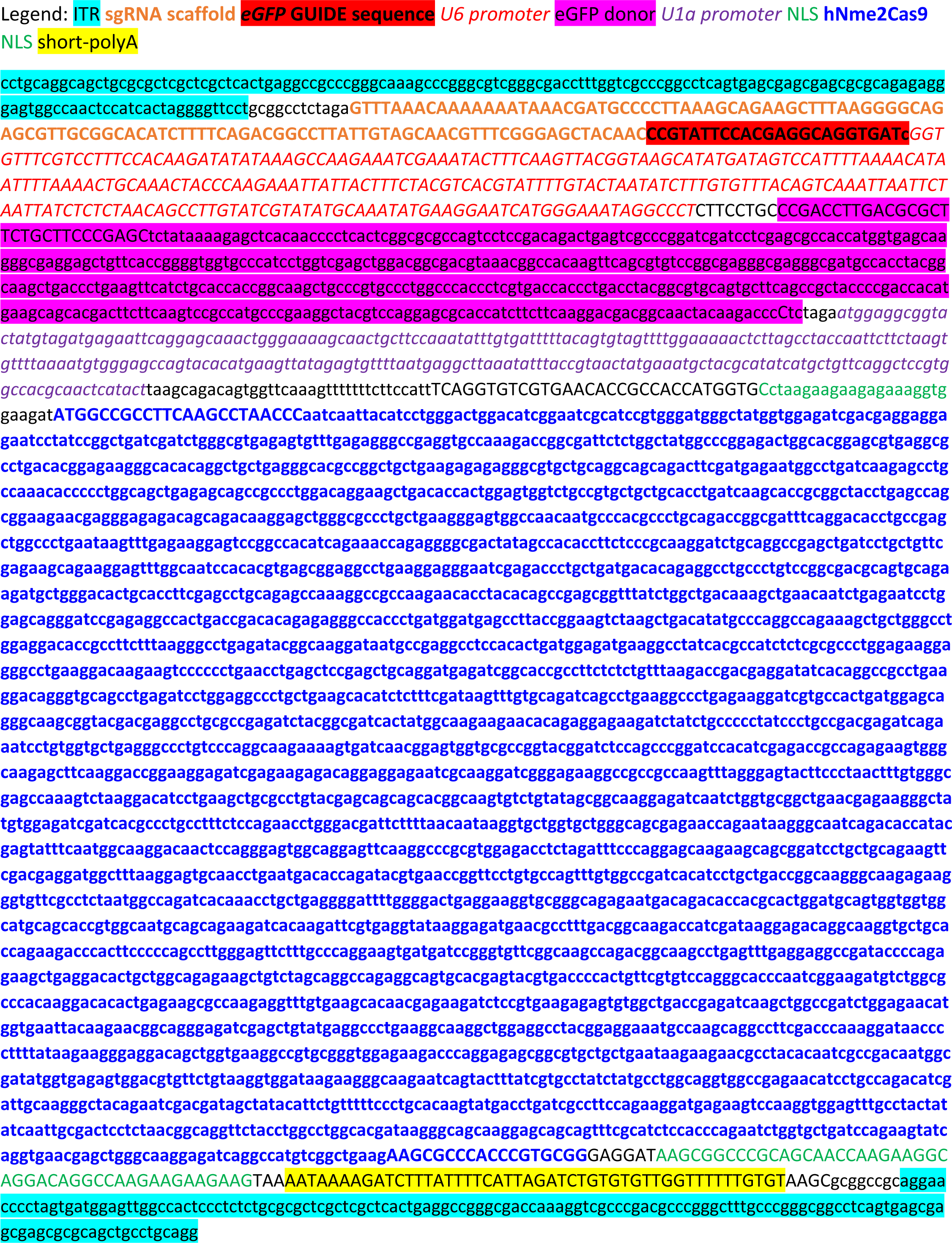

### AAV:DR:cleaved-Design D Plasmid to correct *eGfp* in HEK293T TLR-Multi-Cas-Variant 1 (MCV1)

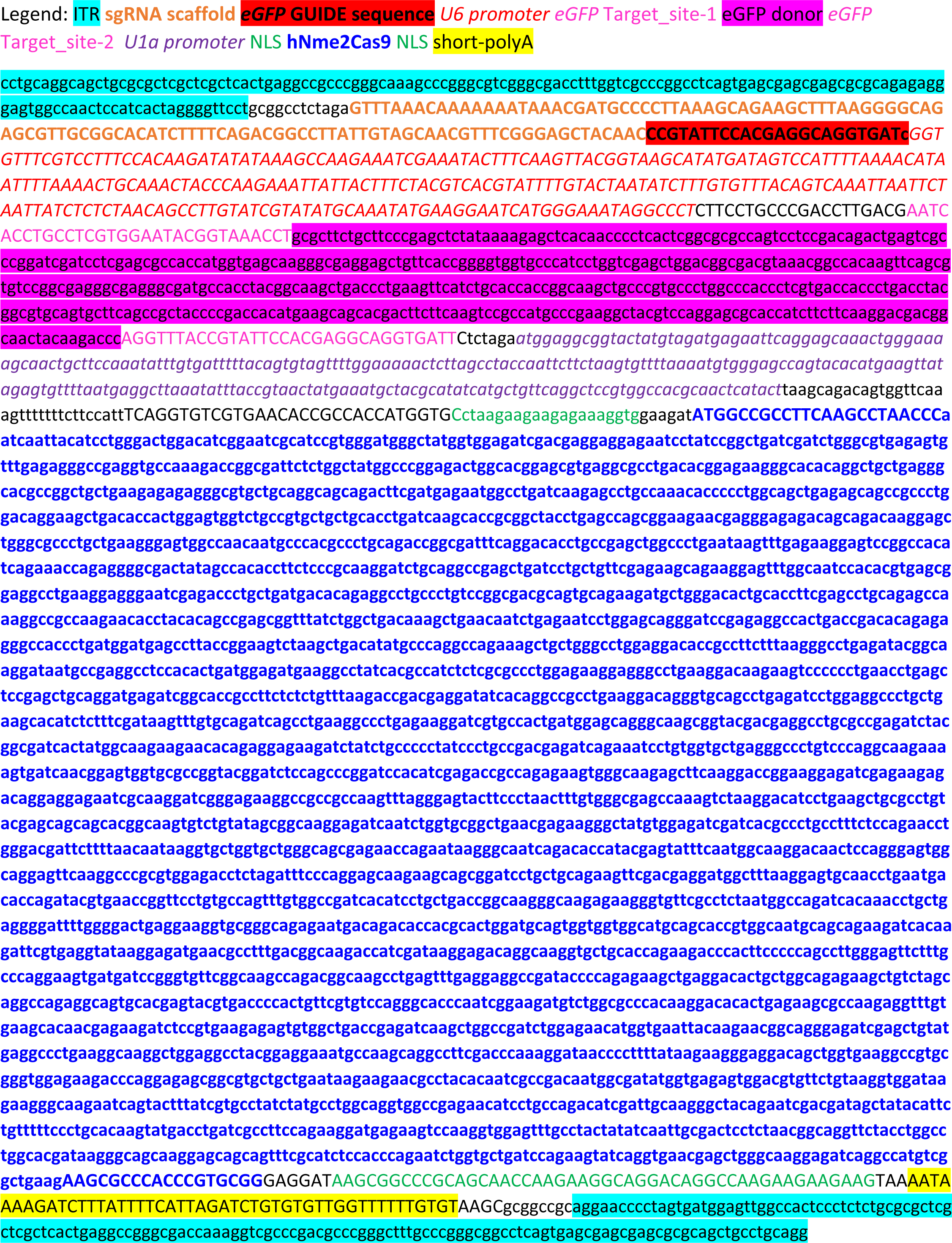

### Figure exemplifying the gating strategy of flow cytometry

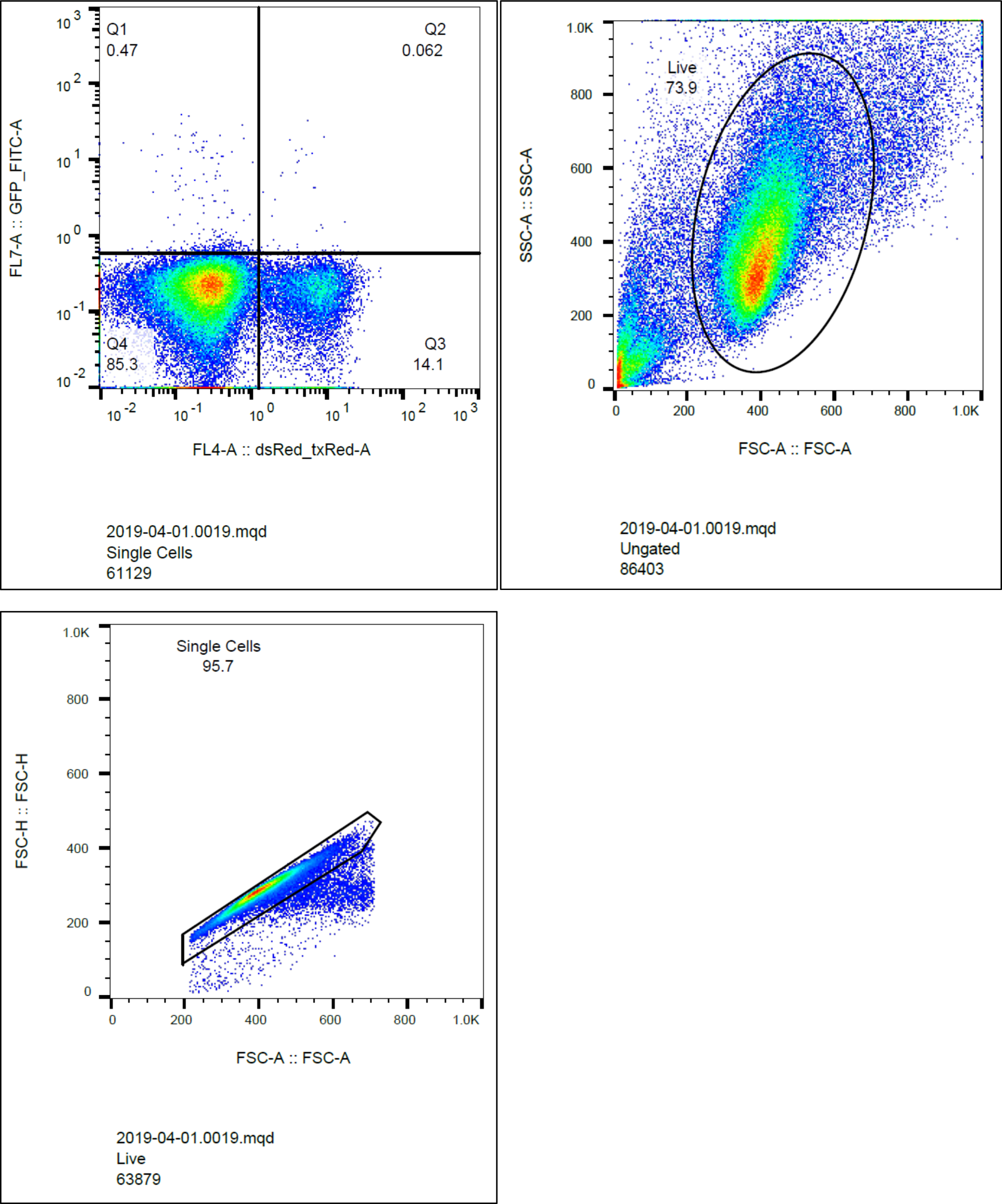

## References

1. Jinek, M. et al. A programmable dual-RNA-guided DNA endonuclease in adaptive bacterial immunity. Science 337, 816–821 (2012).

2. Gasiunas, G., Barrangou, R., Horvath, P. & Siksnys, V. Cas9-crRNA ribonucleoprotein complex mediates specific DNA cleavage for adaptive immunity in bacteria. Proc. Natl. Acad. Sci. U. S. A. 109, E2579–86 (2012).

3. Cong, L. et al. Multiplex genome engineering using CRISPR/Cas systems. Science 339, 819–823 (2013).

4. Mali, P. et al. RNA-guided human genome engineering via Cas9. Science 339, 823–826 (2013).

5. Hwang, W. Y. et al. Efficient genome editing in zebrafish using a CRISPR-Cas system. Nat. Biotechnol. 31, 227–229 (2013).

6. Cho, S. W., Kim, S., Kim, J. M. & Kim, J.-S. Targeted genome engineering in human cells with the Cas9 RNA-guided endonuclease. Nat. Biotechnol. 31, 230–232 (2013).

7. Jiang, W., Bikard, D., Cox, D., Zhang, F. & Marraffini, L. A. RNA-guided editing of bacterial genomes using CRISPR-Cas systems. Nat. Biotechnol. 31, 233–239 (2013).

8. Jinek, M. et al. RNA-programmed genome editing in human cells. Elife 2, e00471 (2013).

9. Zetsche, B. et al. Cpf1 is a single RNA-guided endonuclease of a class 2 CRISPR-Cas system. Cell 163, 759–771 (2015).

10. Liu, J.-J. et al. CasX enzymes comprise a distinct family of RNA-guided genome editors. Nature 566, 218–223 (2019).

11. Pausch, P. et al. CRISPR-CasΦ from huge phages is a hypercompact genome editor. Science 369, 333–337 (2020).

12. Doudna, J. A. The promise and challenge of therapeutic genome editing. Nature 578, 229–236 (2020).

13. Anzalone, A. V., Koblan, L. W. & Liu, D. R. Genome editing with CRISPR-Cas nucleases, base editors, transposases and prime editors. Nat. Biotechnol. 38, 824–844 (2020).

14. Deveau, H. et al. Phage response to CRISPR-encoded resistance in Streptococcus thermophilus. J. Bacteriol. 190, 1390–1400 (2008).

15. Sapranauskas, R. et al. The Streptococcus thermophilus CRISPR/Cas system provides immunity in Escherichia coli. Nucleic Acids Res. 39, 9275–9282 (2011).

16. Yeh, C. D., Richardson, C. D. & Corn, J. E. Advances in genome editing through control of DNA repair pathways. Nat. Cell Biol. 21, 1468–1478 (2019).

17. Rees, H. A. & Liu, D. R. Base editing: precision chemistry on the genome and transcriptome of living cells. Nat. Rev. Genet. 19, 770–788 (2018).

18. Anzalone, A. V. et al. Search-and-replace genome editing without double-strand breaks or donor DNA. Nature 576, 149–157 (2019).

19. Deltcheva, E. et al. CRISPR RNA maturation by trans-encoded small RNA and host factor RNase III. Nature 471, 602–607 (2011).

20. Hou, Z. et al. Efficient genome engineering in human pluripotent stem cells using Cas9 from Neisseria meningitidis. Proc. Natl. Acad. Sci. U. S. A. 110, 15644–15649 (2013).

21. Esvelt, K. M. et al. Orthogonal Cas9 proteins for RNA-guided gene regulation and editing. Nat. Methods 10, 1116–1121 (2013).

22. Ran, F. A. et al. In vivo genome editing using Staphylococcus aureus Cas9. Nature 520, 186–191 (2015).

23. Kim, E. et al. In vivo genome editing with a small Cas9 orthologue derived from Campylobacter jejuni. Nat. Commun. 8, 14500 (2017).

24. Edraki, A. et al. A Compact, High-Accuracy Cas9 with a Dinucleotide PAM for In Vivo Genome Editing. Mol. Cell 73, 714–726.e4 (2019).

25. Hu, Z. et al. A compact Cas9 ortholog from Staphylococcus Auricularis (SauriCas9) expands the DNA targeting scope. PLoS Biol. 18, e3000686 (2020).

26. Wang, D., Zhang, F. & Gao, G. CRISPR-Based Therapeutic Genome Editing: Strategies and In Vivo Delivery by AAV Vectors. Cell 181, 136–150 (2020).

27. Ibraheim, R. et al. All-in-one adeno-associated virus delivery and genome editing by Neisseria meningitidis Cas9 in vivo. Genome Biol. 19, 137 (2018).

28. Agudelo, D. et al. Versatile and robust genome editing with Streptococcus thermophilus CRISPR1-Cas9. Genome Res. 30, 107–117 (2020).

29. Friedland, A. E. et al. Characterization of Staphylococcus aureus Cas9: a smaller Cas9 for all-in-one adeno-associated virus delivery and paired nickase applications. Genome Biol. 16, 257 (2015).

30. Maeder, M. L. et al. Development of a gene-editing approach to restore vision loss in Leber congenital amaurosis type 10. Nat. Med. 25, 229–233 (2019).

31. Chang, Y. J. et al. In vivo multiplex gene targeting with Streptococcus pyogens and Campylobacter jejuni Cas9 for pancreatic cancer modeling in wild-type animal. J. Vet. Sci. 21, e26 (2020).

32. Lee, C. M., Cradick, T. J. & Bao, G. The Neisseria meningitidis CRISPR-Cas9 System Enables Specific Genome Editing in Mammalian Cells. Mol. Ther. 24, 645–654 (2016).

33. Amrani, N. et al. NmeCas9 is an intrinsically high-fidelity genome-editing platform. Genome Biol. 19, 214 (2018).

34. Bisaria, N., Jarmoskaite, I. & Herschlag, D. Lessons from Enzyme Kinetics Reveal Specificity Principles for RNA-Guided Nucleases in RNA Interference and CRISPR-Based Genome Editing. Cell Syst 4, 21–29 (2017).

35. Kleinstiver, B. P. et al. Broadening the targeting range of Staphylococcus aureus CRISPR-Cas9 by modifying PAM recognition. Nat. Biotechnol. 33, 1293–1298 (2015).

36. Shen, M. W. et al. Predictable and precise template-free CRISPR editing of pathogenic variants. Nature 563, 646–651 (2018).

37. Iyer, S. et al. Precise therapeutic gene correction by a simple nuclease-induced double-stranded break. Nature 568, 561–565 (2019).

38. Yang, Y. et al. A dual AAV system enables the Cas9-mediated correction of a metabolic liver disease in newborn mice. Nat. Biotechnol. 34, 334–338 (2016).

39. Nishiyama, J., Mikuni, T. & Yasuda, R. Virus-Mediated Genome Editing via Homology-Directed Repair in Mitotic and Postmitotic Cells in Mammalian Brain. Neuron 96, 755–768.e5 (2017).

40. De Caneva, A. et al. Coupling AAV-mediated promoterless gene targeting to SaCas9 nuclease to efficiently correct liver metabolic diseases. JCI Insight 5, e128863 (2019).

41. Villiger, L. et al. Treatment of a metabolic liver disease by in vivo genome base editing in adult mice. Nat. Med. 24, 1519–1525 (2018).

42. Winter, J. et al. Targeted exon skipping with AAV-mediated split adenine base editors. Cell Discov 5, 41 (2019).

43. Levy, J. M. et al. Cytosine and adenine base editing of the brain, liver, retina, heart and skeletal muscle of mice via adeno-associated viruses. Nat Biomed Eng 4, 97–110 (2020).

44. Yeh, W.-H. et al. In vivo base editing restores sensory transduction and transiently improves auditory function in a mouse model of recessive deafness. Sci. Transl. Med. 12, eaay9101 (2020).

45. Lim, C. K. W. et al. Treatment of a Mouse Model of ALS by In Vivo Base Editing. Mol. Ther. 28, 1177–1189 (2020).

46. Hinderer, C. et al. Severe Toxicity in Nonhuman Primates and Piglets Following High-Dose Intravenous Administration of an Adeno-Associated Virus Vector Expressing Human SMN. Hum. Gene Ther. 29, 285–298 (2018).

47. Wang, L. et al. Meganuclease targeting of PCSK9 in macaque liver leads to stable reduction in serum cholesterol. Nat. Biotechnol. 36, 717–725 (2018).

48. Wilson, J. M. & Flotte, T. R. Moving Forward After Two Deaths in a Gene Therapy Trial of Myotubular Myopathy. Hum. Gene Ther. 31, 695–696 (2020).

49. Krooss, S. A. et al. Ex Vivo/In vivo Gene Editing in Hepatocytes Using ‘All-in-One’ CRISPR-Adeno-Associated Virus Vectors with a Self-Linearizing Repair Template. iScience 23, 100764 (2020).

50. Lee, J. et al. Tissue-restricted genome editing in vivo specified by microRNA-repressible anti-CRISPR proteins. RNA 25, 1421–1431 (2019).

51. Davidson, A. R. et al. Anti-CRISPRs: Protein Inhibitors of CRISPR-Cas Systems. Annu. Rev. Biochem. 89, 309–332 (2020).

52. Marino, N. D., Pinilla-Redondo, R., Csörgő, B. & Bondy-Denomy, J. Anti-CRISPR protein applications: natural brakes for CRISPR-Cas technologies. Nat. Methods 17, 471–479 (2020).

53. Sun, W. et al. Structures of Neisseria meningitidis Cas9 Complexes in Catalytically Poised and Anti-CRISPR-Inhibited States. Mol. Cell 76, 938–952.e5 (2019).

54. Kim, S. et al. CRISPR RNAs trigger innate immune responses in human cells. Genome Res. 28, 367–373 (2018).

55. Hendel, A. et al. Chemically modified guide RNAs enhance CRISPR-Cas genome editing in human primary cells. Nat. Biotechnol. 33, 985–989 (2015).

56. Platt, R. J. et al. CRISPR-Cas9 knockin mice for genome editing and cancer modeling. Cell 159, 440–455 (2014).

57. Geisinger, J. M., Turan, S., Hernandez, S., Spector, L. P. & Calos, M. P. In vivo blunt-end cloning through CRISPR/Cas9-facilitated non-homologous end-joining. Nucleic Acids Res. 44, e76 (2016).

58. Bolukbasi, M. F. et al. Orthogonal Cas9-Cas9 chimeras provide a versatile platform for genome editing. Nat. Commun. 9, 4856 (2018).

59. Guo, T. et al. Harnessing accurate non-homologous end joining for efficient precise deletion in CRISPR/Cas9-mediated genome editing. Genome Biol. 19, 170 (2018).

60. Giannoukos, G. et al. UDiTaSTM, a genome editing detection method for indels and genome rearrangements. BMC Genomics 19, 212 (2018).

61. Xie, J. et al. Short DNA Hairpins Compromise Recombinant Adeno-Associated Virus Genome Homogeneity. Mol. Ther. 25, 1363–1374 (2017).

62. Tai, P. W. L. et al. Adeno-associated Virus Genome Population Sequencing Achieves Full Vector Genome Resolution and Reveals Human-Vector Chimeras. Mol Ther Methods Clin Dev 9, 130–141 (2018).

63. Tran, N. T. et al. AAV-Genome Population Sequencing of Vectors Packaging CRISPR Components Reveals Design-Influenced Heterogeneity. Mol Ther Methods Clin Dev 18, 639–651 (2020).

64. Gao, G. & Sena-Estevez, M. Introducing genes into mammalian cells: viral vectors. in Molecular Cloning: A Laboratory Manual, Vol. 2 (eds. Green, M. R. & Sambrook, J.) 1209–1313 (Cold Spring Harbor Laboratory Press, 2012).

65. Grompe, M. et al. Loss of fumarylacetoacetate hydrolase is responsible for the neonatal hepatic dysfunction phenotype of lethal albino mice. Genes Dev. 7, 2298–2307 (1993).

66. Aponte, J. L. et al. Point mutations in the murine fumarylacetoacetate hydrolase gene: Animal models for the human genetic disorder hereditary tyrosinemia type 1. Proc. Natl. Acad. Sci. U. S. A. 98, 641–645 (2001).

67. Pankowicz, F. P. et al. Reprogramming metabolic pathways in vivo with CRISPR/Cas9 genome editing to treat hereditary tyrosinaemia. Nat. Commun. 7, 12642 (2016).

68. Nelson, C. E. et al. Long-term evaluation of AAV-CRISPR genome editing for Duchenne muscular dystrophy. Nat. Med. 25, 427–432 (2019).

69. Ohmori, T. et al. CRISPR/Cas9-mediated genome editing via postnatal administration of AAV vector cures haemophilia B mice. Sci. Rep. 7, 4159 (2017).

70. Richards, D. Y. et al. AAV-Mediated CRISPR/Cas9 Gene Editing in Murine Phenylketonuria. Mol Ther Methods Clin Dev 17, 234–245 (2020).

71. Gao, J. et al. Viral Vector-Based Delivery of CRISPR/Cas9 and Donor DNA for Homology-Directed Repair in an In Vitro Model for Canine Hemophilia B. Mol. Ther. Nucleic Acids 14, 364–376 (2018).

72. Lee, K. et al. Nanoparticle delivery of Cas9 ribonucleoprotein and donor DNA in vivo induces homology-directed DNA repair. Nat Biomed Eng 1, 889–901 (2017).

73. Shahbazi, R. et al. Targeted homology-directed repair in blood stem and progenitor cells with CRISPR nanoformulations. Nat. Mater. 18, 1124–1132 (2019).

74. Zhang, J.-P. et al. Efficient precise knockin with a double cut HDR donor after CRISPR/Cas9-mediated double-stranded DNA cleavage. Genome Biol. 18, 35 (2017).

75. Petris, G. et al. Hit and go CAS9 delivered through a lentiviral based self-limiting circuit. Nat. Commun. 8, 15334 (2017).

76. Merienne, N. et al. The Self-Inactivating KamiCas9 System for the Editing of CNS Disease Genes. Cell Rep. 20, 2980–2991 (2017).

77. Li, A. et al. A Self-Deleting AAV-CRISPR System for In Vivo Genome Editing. Mol Ther Methods Clin Dev 12, 111–122 (2019).

78. Murphy, K. C. & Campellone, K. G. Lambda Red-mediated recombinogenic engineering of enterohemorrhagic and enteropathogenic E. coli. BMC Mol. Biol. 4, 11 (2003).

79. Lee, J. et al. Potent Cas9 Inhibition in Bacterial and Human Cells by AcrIIC4 and AcrIIC5 Anti-CRISPR Proteins. MBio 9, e02321–18 (2018).

80. Certo, M. T. et al. Tracking genome engineering outcome at individual DNA breakpoints. Nat. Methods 8, 671–676 (2011).

81. Iyer, S. et al. Efficient Homology-directed Repair with Circular ssDNA Donors. bioRxiv 864199 (2019) doi:10.1101/864199.

82. Yin, H. et al. Genome editing with Cas9 in adult mice corrects a disease mutation and phenotype. Nat. Biotechnol. 32, 551–553 (2014).

83. Li, F. et al. Utility of Self-Destructing CRISPR/Cas Constructs for Targeted Gene Editing in the Retina. Hum. Gene Ther. 30, 1349–1360 (2019).

84. Bunge, S. et al. Genotype-phenotype correlations in mucopolysaccharidosis type I using enzyme kinetics, immunoquantification and in vitro turnover studies. Biochim. Biophys. Acta 1407, 249–256 (1998).

85. Palmer, D. J., Turner, D. L. & Ng, P. Production of CRISPR/Cas9-Mediated Self-Cleaving Helper-Dependent Adenoviruses. Mol Ther Methods Clin Dev 13, 432–439 (2019).

86. Wang, D. et al. Characterization of an MPS I-H knock-in mouse that carries a nonsense mutation analogous to the human IDUA-W402X mutation. Mol. Genet. Metab. 99, 62–71 (2010).

87. Hanlon, K. S. et al. High levels of AAV vector integration into CRISPR-induced DNA breaks. Nat. Commun. 10, 4439 (2019).

88. Afgan, E. et al. The Galaxy platform for accessible, reproducible and collaborative biomedical analyses: 2018 update. Nucleic Acids Res. 46, W537–W544 (2018).

89. Edgar, R. C. Search and clustering orders of magnitude faster than BLAST. Bioinformatics 26, 2460–2461 (2010).

90. Wang, D. et al. The designer aminoglycoside NB84 significantly reduces glycosaminoglycan accumulation associated with MPS I-H in the Idua-W392X mouse. Mol. Genet. Metab. 105, 116–125 (2012).

91. Wang, D. et al. Cas9-mediated allelic exchange repairs compound heterozygous recessive mutations in mice. Nat. Biotechnol. 36, 839–842 (2018).

